# Insights into regeneration from the genome, transcriptome and metagenome analysis of *Eisenia fetida*

**DOI:** 10.1101/180612

**Authors:** Aksheev Bhambri, Neeraj Dhaunta, Surendra Singh Patel, Mitali Hardikar, Nagesh Srikakulam, Shruti Shridhar, Shamsudheen Vellarikkal, Hemant Suryawanshi, Rajesh Pandey, Rijith Jayarajan, Ankit Verma, Vikram Kumar, Abhishek Bhatt, Pradeep Gautam, Manish Rai, Jameel Ahmed Khan, Bastian Fromm, Kevin J. Peterson, Vinod Scaria, Sridhar Sivasubbu, Beena Pillai

## Abstract

Earthworms show a wide spectrum of regenerative potential with certain species like *Eisenia fetida* capable of regenerating more than two-thirds of their body while other closely related species, such as *Paranais litoralis* seem to have lost this ability. Earthworms belong to the phylum annelida, in which the genomes of the marine oligochaete *Capitella telata*, and the freshwater leech *Helobdella robusta* have been sequenced and studied. The terrestrial annelids, in spite of their ecological relevance and unique biochemical repertoire, are represented by a single rough genome draft of *Eisenia fetida* (North American isolate), which suggested that extensive duplications have led to a large number of HOX genes in this annelid. Herein, we report the draft genome sequence of *Eisenia fetida* (Indian isolate), a terrestrial redworm widely used for vermicomposting assembled using short reads and mate-pair reads. An in-depth analysis of the miRNome of the worm, showed that many miRNA gene families have also undergone extensive duplications. Genes for several important proteins such as sialidases and neurotrophins were identified by RNA sequencing of tissue samples. We also used *de novo* assembled RNA-Seq data to identify genes that are differentially expressed during regeneration, both in the newly regenerating cells and in the adjacent tissue. Sox4, a master regulator of TGF-beta induced epithelial-mesenchymal transition was induced in the newly regenerated tissue. The regeneration of the ventral nerve cord was also accompanied by the induction of nerve growth factor and neurofilament genes. The metagenome of the worm, characterized using 16S rRNA sequencing, revealed the identity of several bacterial species that reside in the nephridia of the worm. Comparison of the bodywall and cocoon metagenomes showed exclusion of hereditary symbionts in the regenerated tissue. In summary, we present extensive genome, transcriptome and metagenome data to establish the transcriptome and metagenome dynamics during regeneration.

## Introduction

Members of phylum Annelida, commonly represented by earthworms and leeches, occupy a variety of niches from the soils in our gardens to marine sediments and are cosmopolitan in distribution. They sieve our soils^1^, parasitize on a variety of marine and terrestrial hosts^2^, burrow through deep marine sediments^3^ and are also found as fossilized remains from the Cambrian era^4^. Broadly divided into two major classes; Clitellata which is further divided into Oligocheata (earthworms) and Hirudinea (leeches) and the class Polycheata (largely comprising marine worms), the systematics of this group of segmented worms undergoes constant reworking in the light of modern molecular tools employed for classification^5,6^. They belong to the Superphylum Lophotrochozoa, which encompass phyla including the mollusks and flatworms, grouped according to protein coding surveys into a 402-ortholog dataset^7^.

Many earthworm species have a remarkable ability to regenerate part of the body lost due to injury. In the presence of model organisms such as Hydra^8^ or planarians^9^ for studying regeneration, annelids pose an interesting new challenge. There exist large variations in the processes of regenerative capacities within annelids; with leeches which show none to those organisms which can produce an entire individual from a mid-body segment such as some sabellids, chaetopterids and lumbriculids^10^. All this hints towards a diversity of mechanisms for regeneration and asexual reproduction in annelids.

Earthworms also show intriguing behaviors like the ability to distinguish light of different wavelengths^6,11^ and respond to tactile stimuli^12^, vibrations^13^ and dragging objects along directions that offer least resistance^14^. They harbor a large and diverse microbiome in their gut and inherit certain bacteria as part of their nephridial metagenome^15^. They work in close proximity to microbial decomposers to reduce organic matter both depending on microbes and inducing changes in the microbiome favouring some species over others through the production of secondary metabolites and anti-microbial agents^1^.

Superphylum Ecdysozoa, which includes *Caenorhabditis.elegans* (nematodes) and the fruitfly, and the Superphylum Deuterostomia, which includes the vertebrates are represented by several species with completely sequenced genomes^16-18^. These whole-genome sequences have given us new perspectives on the questions of evolution of genomes and the complexities of gene regulatory networks in higher organisms. Genomes of representative terrestrial annelids like the endogeic earthworm, *Lumbricus* sp. and the epigeic vermicomposting worm *Eisenia fetida* can provide a point of comparison with closely related marine oligochaetes *Platynereis dumerilii*^*19*^ and *Capitella teleta* ^*20*^. Comparative genomics of these related species can potentially offer novel insights into our ecosystems, commercially relevant anti-microbials, secondary metabolites and agriculturally important microbial symbionts.

Here, we report the genome sequence of the epigeic vermicomposting worm *Eisenia fetida*, also known as red-worm, along with an extensive analysis of transcriptome dynamics during regeneration. Zwarycz et. al., based on HOX gene analysis from a draft genome have previously reported that the *E. fetida* genome has undergone extensive duplications^21^. We used miRNA as indicators of phylogenetic history to find that like Hox genes, several miRNA families have multiple paralogues in *E. fetida*. In the earthworm metagenome, unique and unculturable microbes were identified. Through extensive annotation of transcriptomic data, we also provide insights into geneic features that underlie their regenerative ability. We report the presence of a neurotrophin gene, with limited similarity to mammalian nerve growth factor family, that is upregulated during regeneration. We show that genes known to enhance neural regeneration are induced during earthworm regeneration implying that conserved molecular pathways are involved in the gene expression program during regeneration.

## Materials and Methods

### DNA and RNA Isolation

*Eisenia fetida* earthworms were procured from farmers engaged in vermicomposting and then maintained in a culture in the laboratory at around 22°C with moderate humidity. The worms were rinsed in tap water to remove any soil attached to the worms. They were then fixed in 70% ethanol for 5 minutes. A platform was used to pin the worm from both ends in order to dissect out the gut. The bodywall was then cleaned thoroughly to remove all residual soil matter. The DNA was then isolated according to a protocol adapted from Adlouni et al^22^. Briefly, the tissue was homogenized using liquid nitrogen and dissolved in 1ml of DNA extraction buffer (NaCl 100mM; EDTA 50mM; Sucrose 7%; SDS 0.5%; Tris base 100mM; pH 8.8). Fifty microlitres of Proteinase K (10mg/ml) was added and the homogenate was incubated at 65°C for 2 hours. The proteins were precipitated by 120uL of 8M Potassium acetate at 4°C. Precipitated proteins were removed by centrifugation at 10,000g while the supernatant was treated with equal volume of Phenol: Chloroform: Isoamyl alcohol on ice. The aqueous layer was recovered after a 15minute spin at 10,000g and DNA was precipitated by equal volume of Isopropanol. DNA was centrifuged at 10,000g and desalted by repeated 70% ethanol washes. The pellet was air dried and dissolved in Tris EDTA buffer (pH 7.5).

Samples from regenerating worms were collected at three different time points - 0 days post cut (dpc), 15dpc, 20dpc and 30dpc. The tissue collected at 0dpc from 60 ± 6 segments was used as reference (0C) for comparison. The regenerated tissue and the tissue adjacent to it, termed control, were collected separately. RNA was isolated using the Trizol method. Briefly, the tissues were frozen in liquid nitrogen, homogenized using a mortar and pestle in the presence of 1ml Trizol reagent and transferred to a microfuge tube. Phase separation was done by adding 200uL chloroform and centrifugation at 10,000g. The aqueous phase was collected and equal volume of isopropanol was added to precipitate RNA. Pellet was collected by centrifugation at 10,000g, followed by repeated 70% ethanol washes. The RNA was then air dried and dissolved in nuclease-free water.

### DNA and RNA sequencing

Approximately, 5ug of DNA, taken from three different worms was sheared using Covaris S220 platform and desired fragment sizes were selected by agarose gel electrophoresis. Paired end libraries of fragment sizes 200bps and 500bps were constructed using Illumina TruSeq DNA Library Prep Kit, while three mate-pair libraries were constructed of insert sizes 10Kb, 7Kb and 5Kb (average sizes since a range was cut from the agarose gel) using Nextra Mate-Pair Library Prep Kit according to manufacturer’s protocol. It should be noted that for the mate-pair library construction, shearing was performed after circularization, as per manufacturer’s guidelines. Briefly, the sheared DNA was end-repaired and purified using AMPure XP beads (Beckman Coulter). This end repaired DNA was then A-tailed and ligated to adapters. The ligated DNA fragments were then amplified by PCR and purified again using AMPure XP beads generating paired end library for sequencing on the Illumina Platform. The 200bp-insert library was sequenced using Illumina GAnalyzer II Platform while the 500bps-insert library was sequenced using HiSeq 2500. For generating long reads (not used in the assembly), Roche 454 libraries were constructed as per manufacturer’s protocol using Shotgun sequencing approach of GS FLX+ system from Roche. One μg of gDNA (estimated using Qubit high sensitivity assay) was used for rapid library preparation which includes DNA fragmentation by nebulization, fragment end repair, adaptor ligation and small fragment removal by AMPure beads based purification. Library was quantified using Quantifluor (Promega) and qualitatively assessed by Bioanalyzer (High sensitivity chip from Agilent). The average fragment length of library was between 1400-1800 bps with <10% of total fragments under 650 bps. This was used to make working aliquots (1 x 10^7^ molecules/μl) for emulsion PCR (emPCR) standardization. For clonal amplification of the library, the optimal amount of DNA titrated using small volume emPCR was used for sequencing. After emPCR optimization, eight DNA copies per bead (cpb) were used for large volume emPCR. The enriched beads were sequenced using pyrosequencing on a picotiter plate. The raw data comprising of series of images were extracted, normalized and converted into flowgrams. These flowgrams are used during signal processing to generate analysis ready sequencing reads. The libraries were sequenced in two region format of the picotiter plate. Lastly, for Oxford Nanopore sequencing, MAP 06 kit was used according to manufacturer’s protocol. Briefly, the DNA was sheared using Covaris g-Tube at 3000rpm, end repaired and A-tailed. Then, ligation was done with hairpin adapters linked to Biotin labelled motor protein. The library was purified using Dynabeads M-280 and then sequenced. Metrichor was used for basecalling. Poretools was used to get fasta sequences^23^.

Approximately, 1ug of RNA was taken per sample and RNA sequencing libraries were made using TruSeq v2 Library Prep Kit as per manufacturer’s protocol. Briefly, the RNA was polyA selected using OligodT magnetic beads followed by shearing into 200-500bp fragments. This sheared RNA was then used to generate cDNA. The cDNA was end-repaired to blunt ends. These blunt ends were then A-tailed i.e. a “A” overhang was added so as to ligate the adapters in the next step. The adapter ligated cDNA was then amplified by PCR and purified by AMPure XP beads. This library was then quality checked and sequenced on Illumina HiSeq 2500. For small RNA sequencing, total RNA was isolated from bodywall and sequenced on the illumina platform. ∼800ng of Total RNA was used as the starting material. Briefly, 3’ adaptors were ligated to the specific 3’OH group of microRNAs followed by 5’ adaptor ligation. The ligated products were reverse transcribed with Superscript III Reverse transcriptase by priming with Reverse transcriptase primers. The cDNA was enriched and barcoded by PCR (15 cycles) and cleaned using Polyacrylamide gel. The library was size selected in the range of 140 – 160 bp followed by overnight gel elution and salt precipitation using Glycogen, 3M Sodium Acetate and absolute ethanol. The precipitate was re-suspended in Nuclease free water. The prepared library was quantified using Qubit Fluorometer, and validated for quality on High Sensitivity Bioanalyzer Chip (Agilent).

### DNA Isolation, sequencing and analysis for metagenomics

The worms were kept in water for a few hours to allow them to purge out gut contents. The gut microbes were avoided by carefully removing the gut before rinsing the bodywall several times to dissociate any residual microbes. DNA was isolated as described above. 16S rDNA PCR was done using specific primers and amplicons were used to generate libraries for Ion Torrent PGM sequencing. Metagenome analysis was performed using MG-RAST ^24^.

### Genome Assembly

The quality of sequencing data was checked using FastQC and reads of phred score>33 were used for further Before starting the assembly, the quality check of the reads was done using FASTQC. The reads were trimmed to remove low quality reads (Phred score 33) using Trimmomatic. Further, reads were filtered for microbial contamination by removing reads matching any organism using an in-house database. The draft genome was made using CLC Genomics Workbench with a word size of 64 and bubble size of 50. The paired end data (obtained from Illumina HiSeq 2500 with an insert size of 500bps) was used for assembling contigs while mate pair data (with an insert size of 3.5-5.5Kb) was used for scaffolding the contigs for the final asesmbly. The resulting assembly was assessed using Assemblathon script to get the N50 statistics.

### Transcriptome Analysis and Annotation

The quality of RNA sequencing reads were checked using FastQC and reads of phred score>33 were used for adapter trimming by Trimmomatic (default parameters). The data was then *de novo* assembled using Trinity package ^25^. The assemblies were annotated using Transdecoder for finding ORFs. The peptides were compared to Uniprot entries using default parameters in BLAST. Differential expression analysis required a complete genome assembled and annotated so as to align reads to the reference. To get a common reference, all the samples were assembled together in a single assembly using Trinity and annotated by Trinotate. This assembly was then used as a reference to align reads of individual samples using Bowtie^26^ and was then assembled using Cufflinks. Read counts were obtained from alignment files using HTSeq^27^. Differential expression profiles were generated using DESeq2^28^ with multiple sample correction done using Benjamini Hochberg. The data generated was then analyzed using the MATLAB suite. Small RNA sequencing data was analyzed using miRminer^29,30^ pipeline. Functional classification of differentially expressed genes was carried out using DAVID. Comparisons were done against user-defined background genelist of Uniprot IDs of *E. fetida* orthologs. Benjamini-Hochberg corrected p-value <10^-4^ was used to select over-represented GO terms.

### RT PCR

The cDNA was prepared using NEB MMULV-RT enzyme using Random Hexamers. RT PCR was performed using Takara Sybr II in Roche LightCycler 480. The primers used have been listed in Supplementary Table (S1). The products were visualized by agarose gel electrophoresis.

### PCR cloning and Sanger sequencing

The cDNA was prepared using Transcriptor High Fidelity cDNA synthesis kit (Roche). The cDNA was used for PCR reaction using Fermentas Taq Polymerase. The PCR product was cloned in pET23A and sequenced using T7 primers in ABI 3130xl Genetic Analyzer according to manufacturer’s protocol.

### Cryosectioning and histology

The earthworm with regenerated tails were fixed in 4% para-formaldehyde solution at room temperature for four to five hours. After fixation, the samples were washed two to three times with phosphate buffered saline (pH 7.0) at room temperature. After, washing, the samples were kept in a 15% sucrose solution (in PBS) and later in 30% sucrose solution at 4°C until the tissue was immersed. The regenerated portion of earthworm was then cut into appropriate sizes to be mounted into cryostat. The samples were freezed at −20°C in the presence of Jung Tissue Freezing Medium^(tm)^. Cross-sections of 20μm thickness were cut from parts of the regenerated portion and immediately transferred onto glass slides coated with gelatin by an artist brush and thaw mounted. The images were taken using a standard brightfield microscope at 5X and 10X magnifications after hematoxylin and eosin staining.

### In situ hybridization

Regenerated earthworms (15dpc, 20dpc and 30dpc) were collected and washed with autoclaved ultrapure type-1 water, followed by overnight fixation at 4°C in 4% (w/v) para-formaldehyde (Sigma-Aldrich) prepared in 1XPBS, pH 7.4. Stringent washes using 0.1% tween-20 in 1XPBS were given to the worms with subsequent storage in 100% methanol at −20°C.

The amplified sequence of SOX4 using specific primers (S1) was cloned in TOPO2 dual promoter vector (Invitrogen). HindIII and EcoRV (New England Biolabs) were used to linearize the cloned plasmid followed by in vitro synthesis of riboprobes using digoxigenin labeled UTPs and RNA polymerase (T7 and SP6) as per the manufacture’s guidelines (Roche). Prior to hybridization, the stored worms were rehydrated in a gradient of 75%, 50%, 25% and 0%(v/v) of methanol in 0.1% tween-20 in 1XPBS. The worms were permeabilized by treatment with 20μg/ml Proteinase k for 45 minutes with subsequent fixation in 4% (w/v) PFA. Hybridization of Earthworms using the sense and antisense probes was performed at 65°C in Hybridization Buffer (50% formamide, 1.3XSSC, 5mM EDTA, 0.2% tween-20, 100μg/ml heparin and 50μg/ml yeast RNA in autoclaved ultrapure type-1 water) followed by stringent washing with TBST solution (0.5M NaCl, 0.1M KCl, 0.1M tris-HCl (pH 7.5) and 0.1% tween-20 in autoclaved ultrapure type-1 water). The earthworms were then incubated in 1:2000 dilution of digoxigenin-alkaline phosphatase fab fragments (Roche) in TBST and 10% FBS at room temperature for 4 hours. The earthworms were washed with TBST and allowed to stain using 100mg/ml Nitro-blue tetrazolium (NBT) and 50 mg/ml 5-bromo-4-chloro-3′-indolyphosphate (BCIP) substrate (Roche) in developing solution (0.1M NaCl, 0.1M Tris-HCl (pH 9.5), 0.05M MgCl_2_, 1% tween-20 in autoclaved ultrapure type-1 water). The reaction was terminated after development of signal by washing earthworms with PBS followed by washing with stop solution (150mM NaCl, 1.2mM EDTA and 50mM MgCl_2_.6H_2_O in autoclaved ultrapure type-1 water with pH adjusted to 7.4). For image acquisition, earthworms were mounted in 2.5% methyl cellulose and imaged under Nikon SMZ800N stereomicroscope at 8X, 6X, 4X and 2X magnification.

### Data Availability

The RNA-Seq raw data and the processed data with fold changes have been deposited in GEO (GSE101310). The small RNA sequencing data of the bodywall used for miRNA annotation is available from SRA (SRP115251)

## Results

### *E. fetida* genome sequence

The diploid genome of a group of *Eisenia fetida* worms from a vermicompost culture maintained in the laboratory was sequenced. We confirmed the species of the worms by sequencing the variable region of cytochrome c oxidase amplified using universal primers recommended for DNA barcoding ^31,32^. The whole-genome shotgun sequencing and de novo assembly was based Illumina short reads from the genome and a mate-pair library of 10 for scaffolding. The best assembly provided 4,63,133 contigs (N50= 967bp) and 3,99,006 scaffolds (N50=9Kb) (Table 1). The scaffolds were obtained by assembling 1.3 billion paired-end Illumina reads (read length= 100 bp (HiSeq 2500)) and 398million reads from a mate pair library of 10Kb. The quality of the assembled genome was verified using BUSCO^33^, wherein 33.1% of the 978 BUSCO groups searched were identified in our genome. By assessing the ability to detect the 4329 EST sequences of *E.fetida* that were previously available in Genbank. 85.8% of the previously known ESTs (3718 of 4329) were identified in our genome assembly. We also carried out a de novo assembly of RNA-Seq reads from the body wall of a pool of *E.fetida* worms using the Trinity package. 94.6% (67491 out of 71341) transcripts in at least one scaffold of the genome. We also carried out RNA-Seq and *de novo* assembly for tissue collected from the regenerating region earthworms during three stages of regeneration (15, 20 and 30 days post cut). A combined assembly of all the RNA-Seq data led to the identification of 40062 genes collectively accounting for 71341 transcript isoforms. The mitochondrial genome sequence was assembled as a single contig of 16kb from an assembly of 1.1million Roche 454 reads (read length = 1000bp) and showed high conservation to the mitochondrial genomes of the deep-dwelling earthworm *Lumbricus terrestris* and the marine oligochaete, *Capitella teleta* (^20,34^, Supplementary information S4). During the course of this study Zwarycz et. al. have also sequenced a North American isolate of *E. fetida*^21^. Our assembly has approximately five fold higher contig N50 and Scaffold N50 (see Table1 for comparison of both assemblies). Zwarcyz et. al. reported, from the analysis of HOX genes, that an exceptionally large number of duplication events have occurred in the earthworm lineage^21^.

**Table 1:**
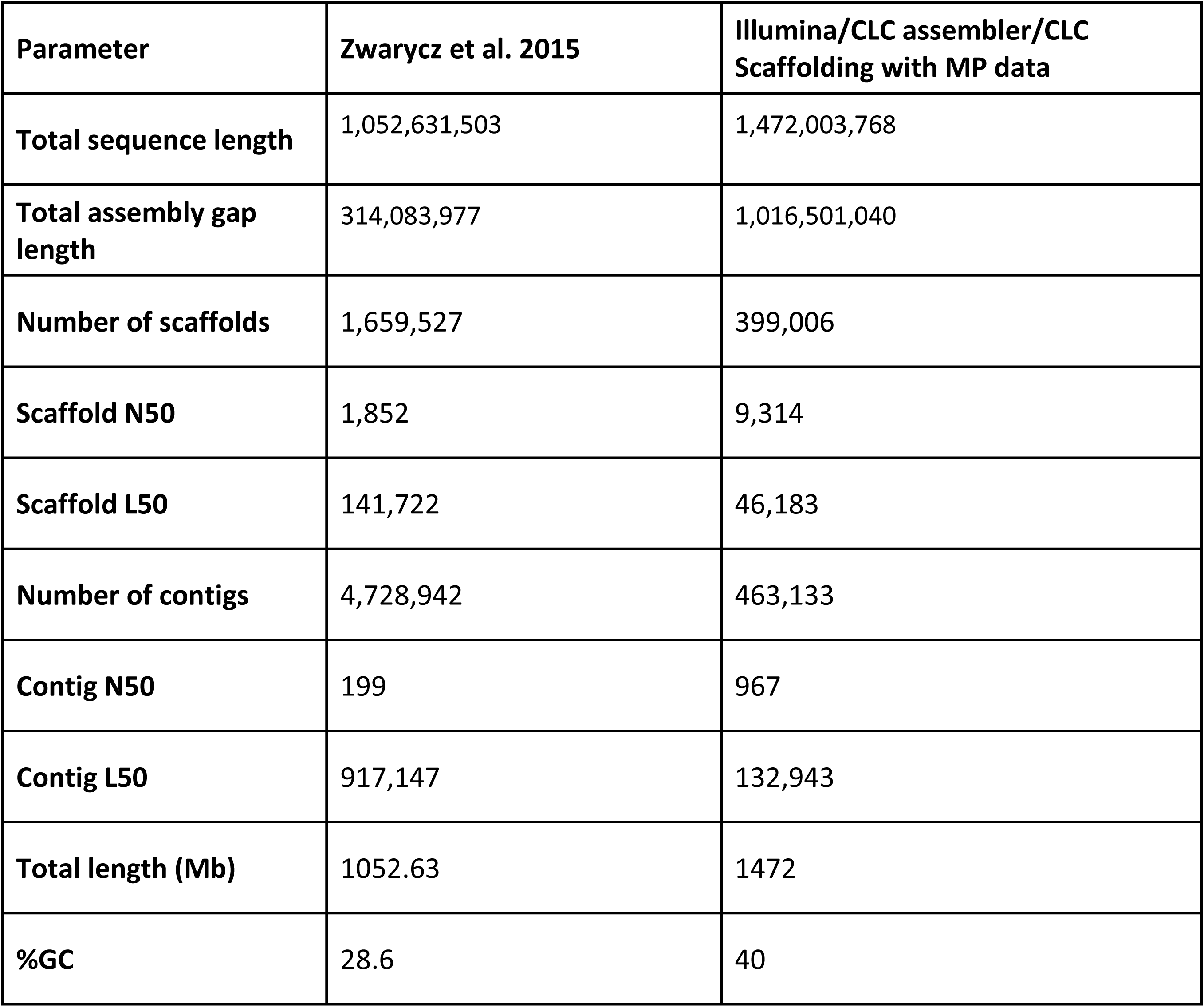
Genome Assembly of *E. fetida*

### miRNome : Genome duplications

Small non-coding RNAs, especially microRNAs have been used as phylogenetic markers to understand evolutionary relations between closely related organisms and genome duplications during evolution. We sequenced the small RNA pool of the *E. fetida* bodywall to identify the microRNAome of the redworm. Using MirMiner^29,30^ we identified 190 microRNA genes belonging to 66 miRNA families, including 18 gene families present only in other lophotrochozoans, neotrochozoans, or annelid worms, and 7 novel miRNA genes not found in *Capitella teleta* or any other metazoan thus far investigated (S2; Fig. 1). Like most metazoan taxa^35,36^, the amount of miRNA family loss appears to be small; these losses, affecting five different miRNA families where neither a locus was detected in the various assemblies nor reads detected in our small RNA library (S2), appear to be shared with *Lumbricus terrestrialis*^30^, and thus most likely occurred after oligochaetes split from *C. teleta*, but before the split between the two earthworm species.

**Figure 1:**
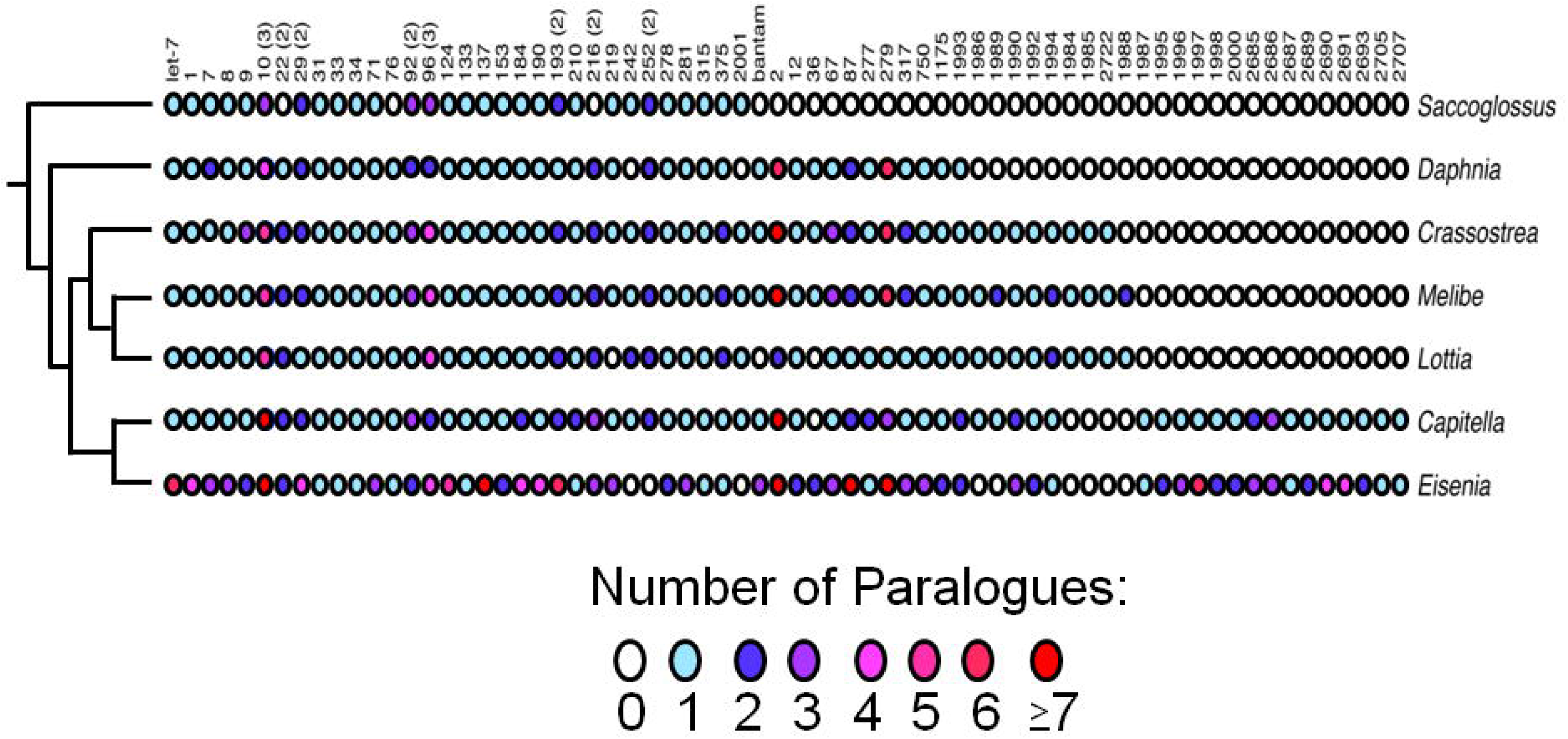
The acquisitional history of 68 microRNA families in select invertebrate bilaterians with well curated microRNAomes. Also shown is the number of microRNA paralogues per family in each taxon. Some families were tandemly duplicated early in bilaterian history including mir-10, mir-22, mir-29 and mir-96; the number of reconstructed genes for each family in the bilaterian last common ancestor is shown in parentheses. Most incidences of paralogy involve these families in these (and most other invertebrate) taxa. *Eisenia* is a notable exception in that most families are characterized by the possession of at least two paralogues, and many families have many more, consistent with the hypothesis that a (or multiple) genome duplication events occurred in this lineage after the split from the polychaete *Capitella.* See S3 for further details.

In contrast to *C. teleta* or any other invertebrate yet investigated, *E. fetida* is rich in miRNA paralogues averaging nearly 2.5 paralogues per ancestral gene (S3). Some families, like Mir-10, Mir-124 and Mir-137, have more copies per gene family than any other animal (including human) yet investigated. Because each of these paralogues is characterized by a unique star (and often mature) sequence, and reads of the mature (and often star) sequence were detected in our small RNA library (S2), the unusually high number of paralogues cannot simply be due to unrecognized heterozygosity (see Zwarycz et al. 2015^21^). Instead, our observation is consistent with what is known regarding homeobox genes in this worm; Zwarycz et al.^21^ found that *E. fetida* has at least 364 homeobox genes with multiple representatives of *Hox* genes including four *Hox1*/*labial*, *Post1* and *Post2* genes; and at least three each of *Hox3*, *Hox5*/*Scr*, *Lox2*, and *Lox4*. Indeed, we found multiple copies of Mir-10 genes associated within the *Hox* cluster^37^ including two mir-10s, three mir-993 genes, two copies of mir-10b (a mir-10 paralogue found in annelids and mollsucs), and one copy each of mir-1991 and mir-10c (an annelid-specific mir-10 paralogue) (S2). Therefore, the microRNAs we found in this worm are consistent with the idea that large, if not whole-scale, gene/genome duplication events occurred in this lineage after its split from *C. teleta*^*21*^, duplications that affected the entire genome including the miRNAs.

### Regeneration in E. fetida

Mature *E. fetida* has 100±10 body segments that show a remarkable potential to regenerate towards the posterior. Based on early studies on *E. fetida*^*38,39*^, a review on the regenerative potential of annelids^40^ mentions that the posterior can also regenerate the anterior segments. However, a more recent report by Nengwen et. al. shows that the posterior, can survive and regenerate the anterior segments, only when the loss is restricted to the first 7 segments^41^. This is in agreement with our own observations which prompted us to focus on the ability of *E. fetida* to regenerate posterior segments. To identify genes differentially expressed during regeneration, we carried out RNA sequencing and created a temporal profile of regeneration in *E. fetida*.

In this study, earthworms were cut transversely at 60 ± 6 segments and allowed to regenerate for 15, 20 or 30 days (Fig.2A). At 15, 20 and 30 days post cut (dpc), the regenerated tissue (henceforth referred to as 15R, 20R and 30R) and the (old) adjacent tissue (henceforth referred to as 15C, 20C and 30C) were collected (Fig.2B) from each batch consisting of 50-100 worms. In one batch, immediately after cutting, tissues were collected from the site (60 ± 6; henceforth called 0C) to be used as a reference for comparing the regenerated and control tissues. Besides these samples, we also carried out RNA-Seq of the posterior (segments 60-100; called P0 for Posterior at 0days) as shown in Fig 2A. Direct comparison of the regenerating tissue with P0 allowed us to establish the extent to which regeneration is completed by 30dpc. Three biological replicates, starting with different batches of earthworms were performed. Total RNA was used for RNA-Seq analysis and the reads were assembled, *de novo*, using Trinity ^25^ and annotated (see Methods). Finally, comparative analysis was performed using DESeq2^28^ to identify genes that were differentially expressed in any of the tissues (15C, 20C, 30C, 15R, 20R or 30R) when compared to the reference (0C). In the tissue adjacent to the site of injury, less than hundred genes (Table 2) were significantly differentially expressed at any point during regeneration. Gene Ontology classification using DAVID also failed to cluster these genes into any specific functional category (Table 3) on the basis of biological process or molecular function (Benjamini Hochberg corrected pVal < 10^-4^). In contrast, in the regenerating tissue, thousands of transcripts from a total of 61,807 transcripts (including isoforms) were differentially expressed (Table 3). The largest number of differentially expressed genes were in the regenerated region at 15dpc. The number of differentially expressed genes steadily reduced as regeneration proceeds at later time points (20dpc and 30dpc). Next, we identified biological processes and molecular functions that were over-represented in the up-regulated or down-regulated genes (Fig 2C-F). Overall, it is clear that in the early stages of regeneration, genes involved in translation, DNA replication and cell division are upregulated while in later stages, ECM remodeling is undertaken.

**Table 2:** Differentially expressed genes from DESeq2 analysis

| **Sample name** | **Number of Differentially Expressed Genes (n=3; Adjusted pVal<0.05)** | **Number of Upregulated Genes (n=3; Adjusted pVal<0.05)** | **Number of Downregulated Genes (n=3; Adjusted pVal<0.05)** |
| --- | --- | --- | --- |
| Segments 60 ± 6 at 0dpc | Reference | Reference | Reference |
| Control at 15dpc (15C) | 75 | 64 | 11 |
| Control at 20dpc (20C) | 72 | 64 | 8 |
| Control at 30dpc (30C) | 63 | 39 | 24 |
| Regenerated at 15dpc (15R) | 8713 | 3589 | 5124 |
| Regenerated at 20dpc (20R) | 5499 | 1887 | 3612 |
| Regenerated at 30dpc (30R) | 1231 | 617 | 614 |
| Posterior segments 60-100 (P0) at 0dpc | 21 | 12 | 9 |

**Table 3:** Gene Ontology classification of Differentially expressed genes.

|  |  | DESeq2 |  |
| --- | --- | --- | --- |
| Sample name |  | UP | DN |
| Control at 15dpc(15C), 20dpc(20C), 30dpc(30C) |  | "n.s" | "n.s" |
| Regenerated at 15dpc (15R) | Biological Process | Mitochondrial translational elongation(24), Translation(79), Mitochondrial translational initiation(14), DNA replication initiation(21), Mitotic nuclear division(83), Mitochondrial translation(21) | Cilium assembly(66), Cell projection organization(44), Cilium movement(20) |
|  | Molecular Function | Structural constituent of ribosome(76), Poly(a) rna binding(197), RNA binding(180), Peptidyl-prolyl cis-trans isomerase activity(22) | Cilium morphogenesis(72), Calcium ion binding(176), Steroid hydroxylase activity(17), Aromatase activity(20), Heme binding(53) |
| Regenerated at 20dpc (20R) | Biological Process | Cell division(72), DNA replication initiation(16), Mitotic nuclear division(53) | Axoneme assembly(14) |
|  | Molecular Function | Extracellular matrix structural constituent(25) | Protein homodimerization activity(110), Calcium ion binding(134), Steroid hydroxylase activity(15), Heme binding(44), Aromatase activity(17) |
| Regenerated at 30dpc (30R) | Biological Process | Extracellular matrix structural constituent(21), Calcium ion binding(43) | "n.s" |
|  | Molecular Function | Extracellular matrix organization(20) | Steroid hydroxylase activity(8), Oxidoreductase activity(7) |

Only GO categories with Benjamini Hochberg adjusted pVAL<10<sup>-4</sup> have been included. No significant classes found ="n.s"; Number of genes in parantheses.

**Figure 2:**
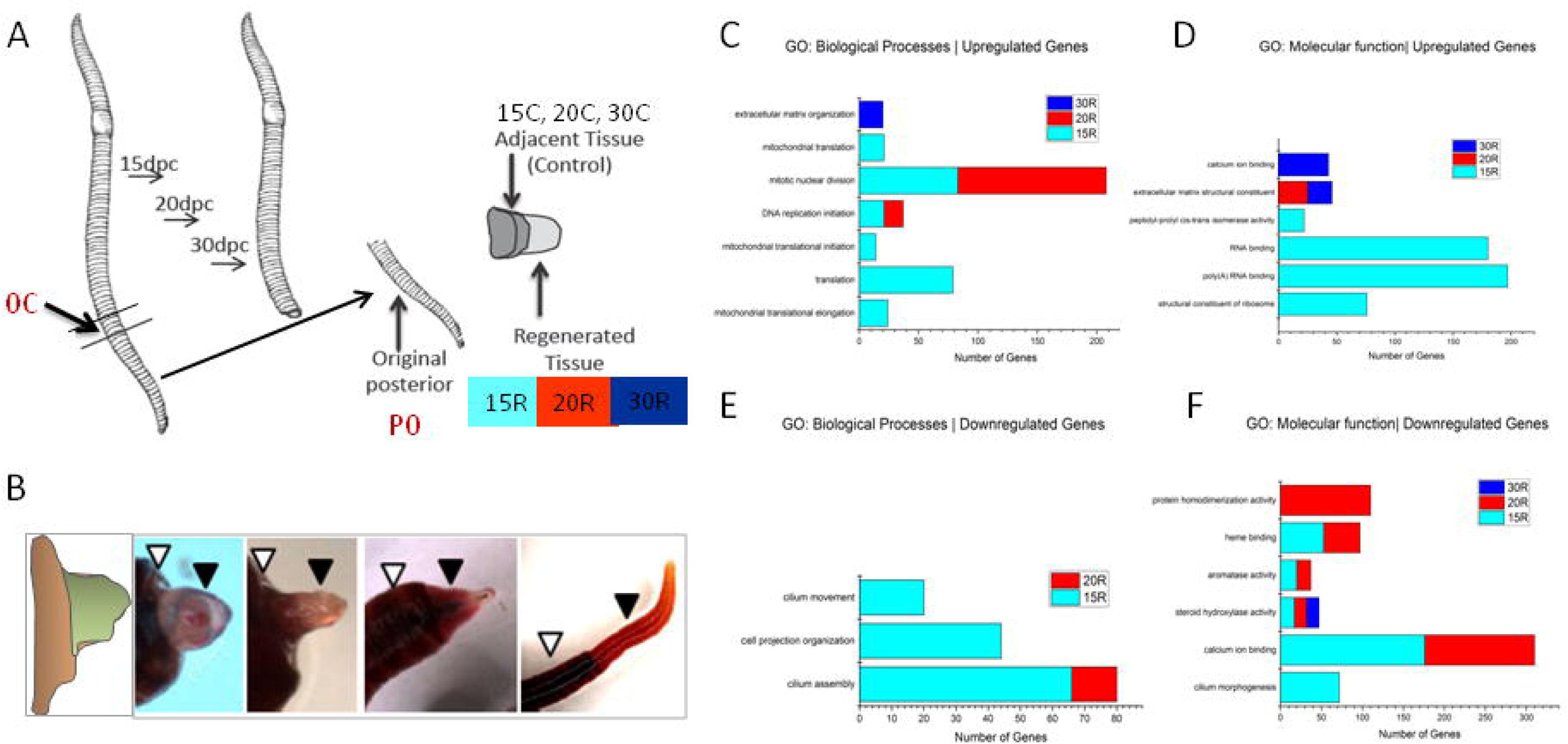
Functional classification of differentially expressed genes in the regenerating earthworm. (A) Schematic representation of the experiment: Earthworms were cut (transverse) at 60 ± 6 segments from the anterior end and anterior portion was allowed to regenerate for 15, 20 or 30 days. At 15, 20 and 30 days post cutting (dpc), the regenerated tissue and the (old) adjacent control tissue was collected. Total RNA was used for RNA-Seq analysis. (B) A typical worm during various stages of regeneration showing the regenerating tissue (filled arrowhead) and the adjacent tissue (open arrowhead). Gene ontology classification by DAVID revealed GO terms over-represented in the (C-D) upregulated genes and (E-F) downregulated genes (Benjamini Hochberg adjusted pVal < 10^-4^)

We also compared the changes occurring in the regenerating tissue with the adjacent tissue. Majority of the differentially expressed genes were downregulated in the regenerating tissue when compared to the adjacent tissue (Fig 3A-F). This apparent downregulation of genes maybe a reflection of reduced heterogeneity amongst the undifferentiated, rapidly proliferating cells in the regenerating tissue. The regenerating tissue, initially, is a homogenous mass of undifferentiated cells. Many genes appear downregulated in the regenerating tissue, because the corresponding cell type has not been formed. For instance, cilium associated genes (Fig 2E-F) are down-regulated until the nephridia and sensory cells in the epidermis are restored. The giant extracellular hemoglobin of annelids have been studied extensively, for its exceptional oxygen carrier properties. Interestingly, the expression of the gene for giant extracellular hemoglobin (Fig 3D-F) of *E. fetida* is induced in the regenerating tissue at 15dpc and 20dpc (Adjusted pVal<0.05). By 30dpc the regenerated tissue has fewer differentially expressed genes, suggesting that the program of regeneration is restoring the expression of genes. Our results suggest that, following injury, the regenerating tissue mounts a well-coordinated program of gene expression involving developmental regulators like SOX2 and SOX4.

**Figure 3:**
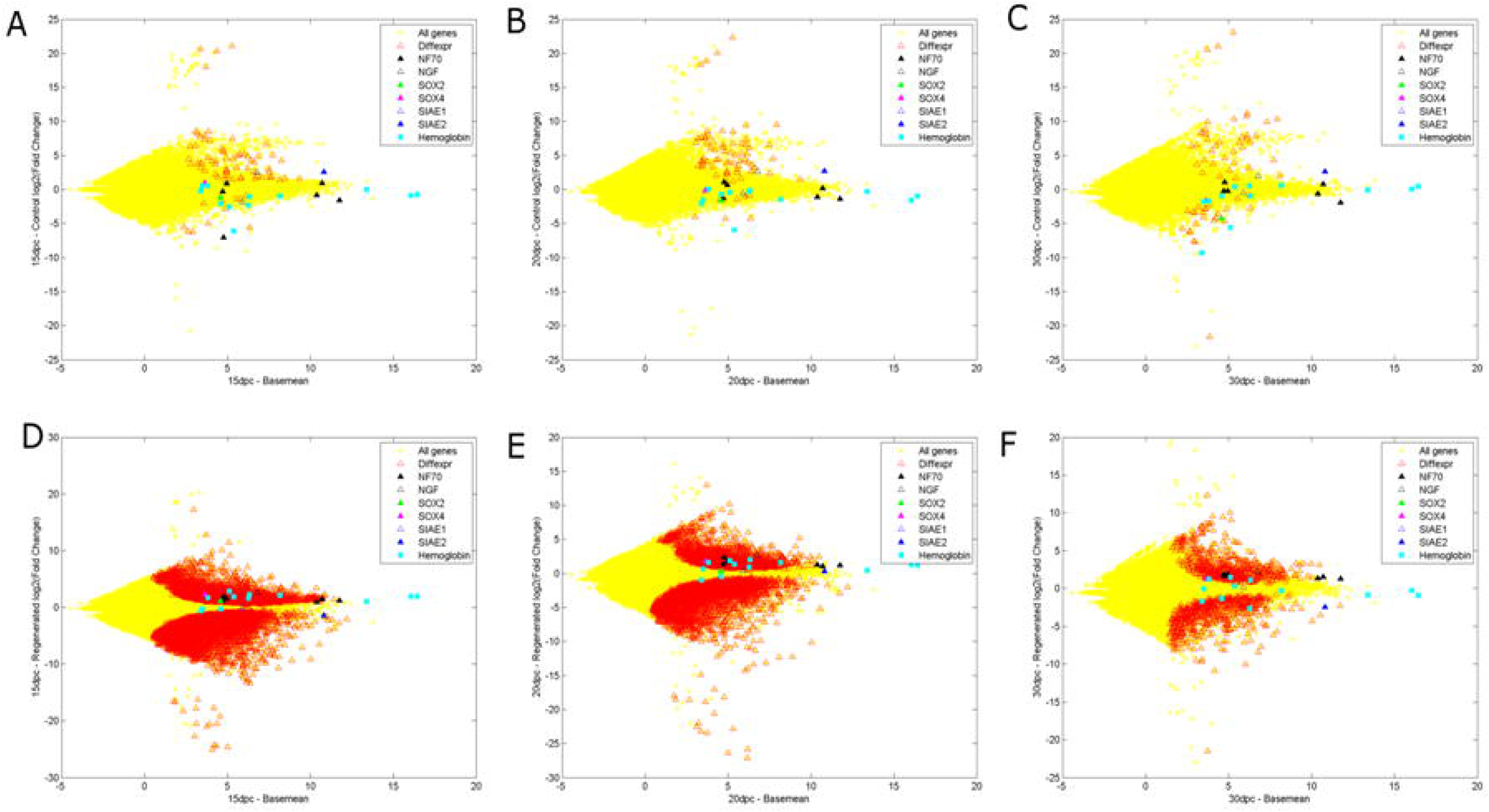
Scatter plot of fold change of genes in the (A-C) control region (Y axis) and (D-F) regenerated region, compared to the basal expression level (basemean; X axis) at 15, 20 and 30 days post cutting. Differentially expressed genes are marked in red. The genes analyzed in detail are highlighted. Divergent expression of two Sialidase isoforms are shown by blue triangles. Neurofilament, NF70 (Black filled triangle) are unaffected in control tissue but induced in the regenerating tissue. Giant extra-cellular hemoglobin genes (cyan Square), Sox4 (pink triangle), Sox2 (green triangle) re-gain expression in the regenerating tissue as regeneration proceeds.

### Regeneration factors

We used k-means clustering to identify co-regulated genes that were upregulated in the regenerating tissue. This cluster is likely to include master regulators of regeneration, since they are highly induced in the regenerating tissue. Interestingly, the most highly induced gene (>100 fold) in the regenerating tissue is Brachyury (Fig. 4A). Brachyury, literally meaning short-tail after the heritable shortened tail of mice that carry a mutation in this transcription factor^42,43^, is expressed in the mesoderm and is required for the specification of posterior structures in fruit flies^44^, sea urchins^45^, nematodes^46^, zebrafish^47^, frogs^48^, chicken^49^ and humans, suggesting a conserved role across evolution ^50,51^. In the hydra and newt, it has been shown to be induced in regenerating tissues^52,53^. Using the Brachyury profile as a reference, we identified a cluster of 913 genes with a strong profile match to Brachyury (Fig. 4B). This cluster consists of several developmental regulators like FGF, BMP2 and WNT signaling genes (Fig. 4C). Another gene, strongly induced in the regenerating posterior is the earthworm ortholog of the homeobox segmentation gene even-skipped (Fig. 4C). In situ hybridization studies in *Capitella telata*^54^ and *Platynereis dumerilii*^55^ have shown that even skipped is a marker of posterior development. Further, it has been implicated in zebrafish fin regeneration^56^ In earthworm, 38 novel genes with no apparent homology in vertebrate genomes are part of the cluster of 913 genes co-expressed with Brachyury during regeneration (Fig.4B-C). In the absence of functional conservation, co-expression of genes can be useful in predicting function, in this instance, implicating these novel genes in regeneration and posterior development.

**Figure 4:**
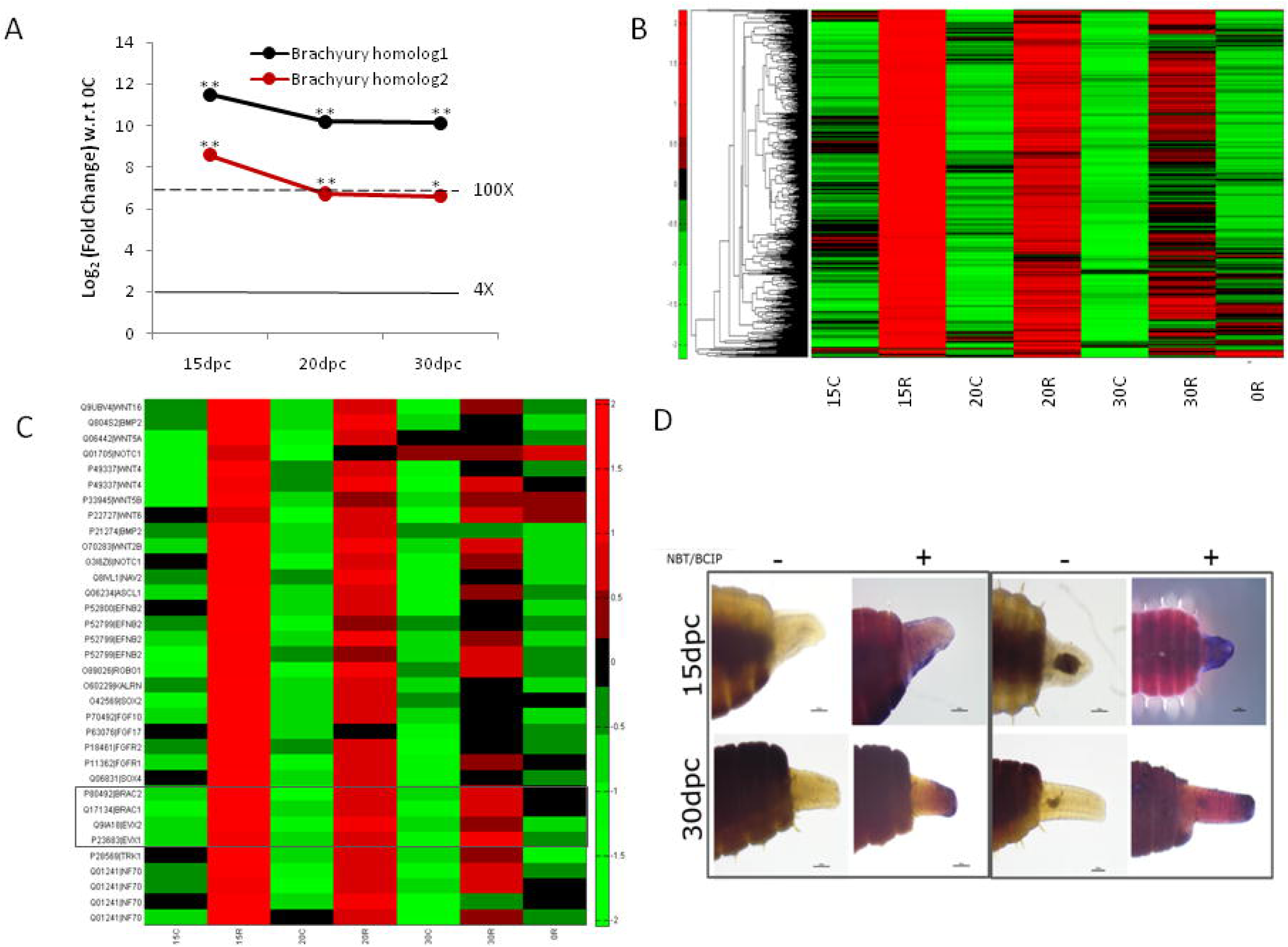
Genes induced in regenerating tissue of *E. fetida*. (A) Brachyury is the most highly induced gene in the regenerating tissue. The two *E. fetida* homologs of *Branchiostoma floridae* (lancelet) Brachyury gene are shown in black and red. *Adjusted pval<0.05; **Adjusted pval<0.005. (B) Cluster of 951 genes that match the profile of Brachyury in k-means clustering (C) Expression pattern of developmental genes that are highly induced in the regenerating tissue. Grey box shows Brachyury and Even-skipped. (D) SOX4 is induced in regenerating tissue at 15dpc and 30dpc. Control (probed with sense probe; left panels 1 and 2) and anti-sense probe (right panels) before (-) and after (+) addition of chromogenic agent (NBT/BCIP).

Sox4 is known to be a master regulator of epithelial to mesenchymal transition^57^. Our Gene Ontology analysis had shown that ECM genes were over-represented amongst the genes upregulated in the regenerating tissue. Therefore, we verified the expression of SOX4, in the regenerating tissue using in situ hybridization. The tip of the regenerating tissue showed a strong and consistent induction of SOX4 (Fig 4D). Taken together, the induction of SOX4, a master regulator of EMT and ECM genes suggest that signalling through the extra-cellular matrix is a vital component of regeneration in *E. fetida*.

### Regeneration of blood and immune cells

A closer examination of the global gene expression profile, especially the genes that are strongly repressed in the regenerating tissue at 15dpc revealed that a group of apparently co-regulated globins were initially depleted in the regenerating tissue. As the newly formed tissue matured, the globin genes became more abundant and eventually at 30dpc were comparable in expression to the adjacent tissue. We believe this is a reflection of the vascularization of the regenerating tissue and the associated presence of blood cells.

Sialidase (SIAE) showed two paralogs with divergent expression pattern at all time points in the regenerated tissue, compared to the adjacent tissue. Sialic acids are the N- or O-substituted derivatives of the 9-carbon monosaccharide^58^, neuraminic acid, N-acetylneuraminic acid being the predominant form in mammalian neural cells while O-acetyl neuraminic acid is a marker for differentiation of immune cells^59,60^. In mammalian genomes, lysosomal and cytosolic isoforms of sialic acid 9-O-acetylesterase catalyze the removal of 9-O-acetylation and play an important role in auto-immunity and B-cell differentiation ^61-63^. Earthworms, especially *E. fetida* have been studied extensively as a model for invertebrate innate immunity ^64^, but the role of SIAE has never been reported. The SIAE gene had no known ortholog in the *C. elegans* and *Drosophila melanogaster* genomes, but genomic analysis revealed the presence of two closely related sialidase genes in the earthworm genome, occuring on two distinct contigs (Fig. 5A). We were intrigued by the divergent expression patterns of the two orthologs, both of which resemble the single human SIAE gene (Fig. 5B). We designed ortholog specific exon-spanning primers and validated our findings from genome assembly and RNA-Seq by RT-PCR. SIAE1, was upregulated in the regenerating tissue by 6.3 fold (data not shown) at 15days post cut while SIAE2 was downregulated (Fig. 5C). The expansion of Hox genes, miRNA genes and the SIAE genes suggest that gene duplication is a recurrent phenomenon in the evolutionary history of *E. fetida*. While the impact of this gene duplication and divergent regulation is yet unknown, the knowledge of novel orthologs may provide deeper insights into aspects of sialic acid mediated immune cell differentiation.

**Figure 5:**
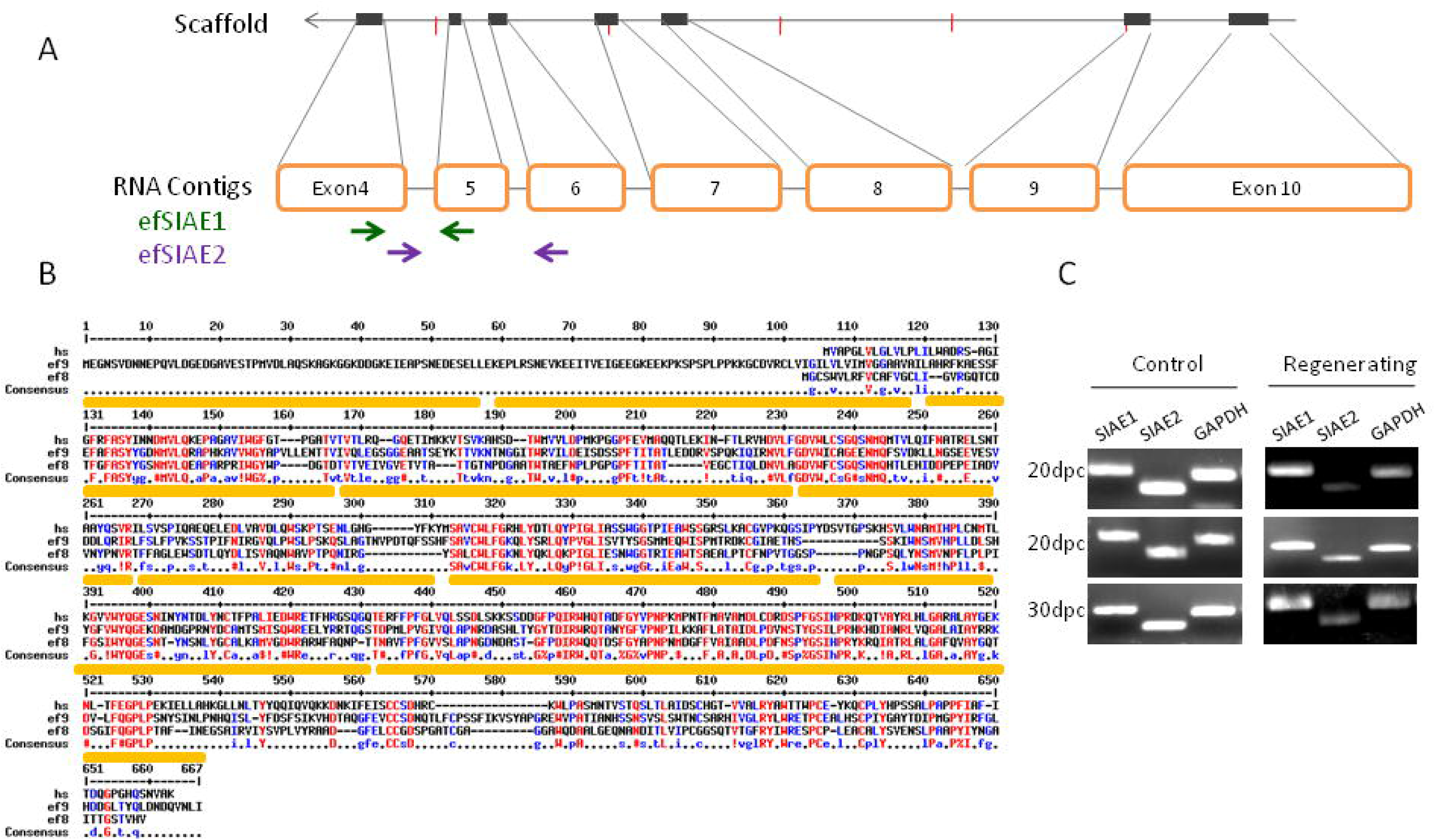
Divergent expression of O-sialic acid esterase homologs during regeneration of *E.fetida*: (A) Gene structure of efSIAE1. Paralog specific primers for efSIAE1 and efSIAE2 are shown by green and purple arrows respectively. (B) Two contigs derived from de novo asssembly of RNA-Seq data showed homology at protein level to O-sialic acid esterase gene of human (hs). One of them was also identified in a scaffold assembled in the genome sequence assembly. (C) RT-PCR using gene specific primers confirmed the divergent expression of the transcripts of efSIAE1 and efSIAE2 in the RNA-Seq data.

### Neural regeneration

Neural regeneration is diminished even in tail regeneration models like the leopard gecko, *E. macularius*, which shows robust regeneration of a hollow surrounding cartilaginous cone devoid of the spinal cord within ^65^. We studied the genes differentially expressed during earthworm regeneration to identify expression profiles associated with neural regeneration in the posterior segments. The strong induction of Nerve Growth Factor and the neurofilament gene, NF70 indicated ongoing neurogenesis (Fig6A,B). Neurofilaments, a major component of the neuronal cytoskeleton are primarily responsible for providing structural support for the axon and regulating axon diameter 66,67. Mammalian neural regeneration is accompanied by the tight regulation and time dependent differential expression of neurofilament genes^68^. Earthworm anatomy shows a ventral nerve chord that connects the dorsal ganglia present in the anterior segments to the rest of the body^69,70^. In agreement with early reports^71^, the ventral cord was visible in the cross-section of the regenerated tissue thus offering a model for nerve regeneration (Fig. 6C).

**Figure 6:**
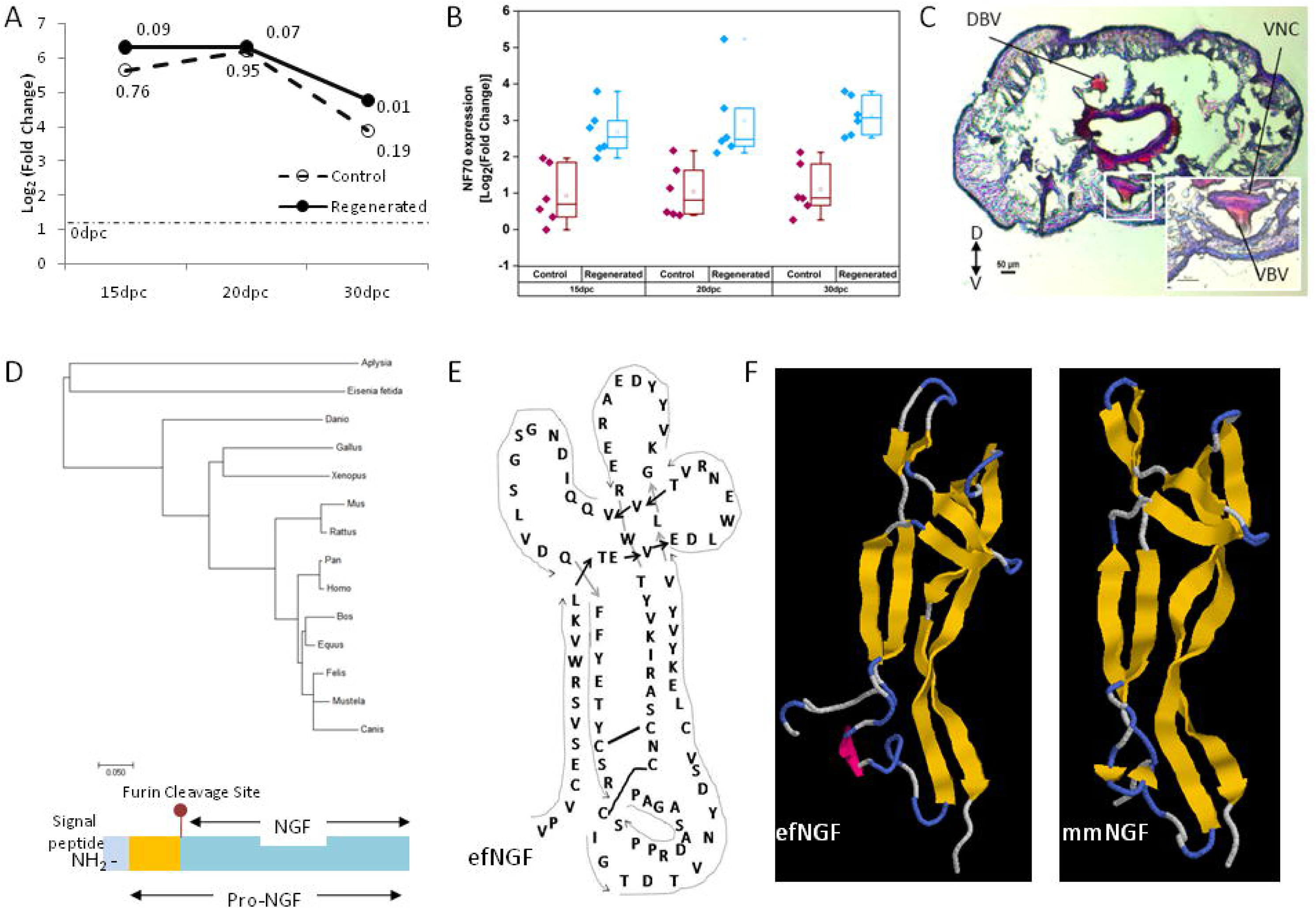
(A) Nerve Growth Factor gene is induced in regenerated tissue (solid black line) at 15dpc, 20dpc and 30dpc. Adjusted pval is mentioned at each point. The gene is also induced in the control tissue but was not statistically significant (dashed line). Stippled line shows the fold change in the posterior (P0) to the control tissue (0C) at the time of injury (0dpc). (B) Neurofilament (NF70) genes are unregulated during regeneration in *E. fetida*. Expression level of various isoforms of NF70 at 15, 20 and 30 days post cutting in the regenerated tissue (15R, 20R, 30R) compared to adjacent control tissue (15C, 20C, 30C). (C) Hematoxylin and Eosin (H&E) staining of cross-section of regenerating earthworm *Eisenia fetida* 20 dpc. (D) Schematic figure of NGF consisting of the signal peptide, pre-NGF and NGF separated by a conserved Furin cleavage site (sequence details in S5) with phylogenetic analysis shows that the *Eisenia fetida* does not resemble other invertebrate NGFs like the molusc NGF in primary protein sequence. (E-F) The signal peptide and the Pro region show minimal homology but the conserved cysteines in the mature NGF reveal structural homology to mouseNGF (mmNGF).

In vertebrates, nerve growth factors play an important role in neurogenesis, axon guidance^72^ and appropriate innervations of regenerated tissue following injury. Nerve growth factors were not known in invertebrates and was even contested until recently. The presence of a nerve growth factor in *Aplysia* was thought to be a unique feature of molluscs amongst invertebrates ^73^. The regenerative ability of *E. fetida* prompted us to explore the newly sequenced genome for NGF-like genes. We report here, the identification of a 747bp pro-NGF which can give rise to a 681bp mature form with limited similarity to mammalian NGF. The conserved gene architecture with the presence of the characteristic furin cleavage site allowed us to align invertebrate and vertebrate NGF like genes (Fig. 6D-F), in spite of the limited homology (Supplementary Information S5). The annotation of the *E. fetida* transcriptome also comprises TrkB receptors, known to bind nerve growth factors. As shown in Figure 4C, TRK1, encoding TrkB, was amongst the highly induced genes in the regenerating tissue.

### Metagenome

Earthworms exist in close proximity to microbes, working symbiotically with a complex metagenome to strongly impact the environment^1^. The genome of the earthworm would be incomplete without knowledge of the microbes that reside in its tissues as obligate symbionts and are inherited through cocoons. Besides cocoons, we carried out metagenome sequencing of the normal and regenerating bodywall. It is known that phagocytic cells of *E. fetida* called coelomocytes, engulf bacteria and form structures called brown bodies that are carried in the coelom and eventually excreted ^74,75^. In spite of the extensive rinsing of the bodywall, we cannot rule out the possibility that some of these microbes are present in the coelomic cavity. To assess the extent of vertical transmission of the metagenome we carried out a comparative metagenomics of the cocoon, and found that a large fraction of species were common between the adult bodywall and the cocoon (Fig. 7A). Currently, it is known that *Verminephrobacter eiseniae* colonizes the excretory structures, called nephridia, found in the earthworm bodywall ^76^. Another uncultivable bacterial species found in tripartite symbiosis with the earthworm and *Verminephrobacter* sp. is the *Flexibacter*, proposed to be renamed to *Candidatus Nephrothrix* (Moller et al., 2015). In agreement with the existing reports, we found both these organisms in the normal earthworm bodywall. Besides these extracellular symbionts, the earthworm bodywall harbors a complex metagenome consisting of diverse microbes from several eubacterial and archaebacterial classes such as Corynebacteria, Flavobacteria and Propionibacteria (Fig. 7A). Our findings suggest that vertical transmission of microbes is a widespread phenomenon in *E. fetida* and may extend to some species that are not classified currently.

**Figure 7:**
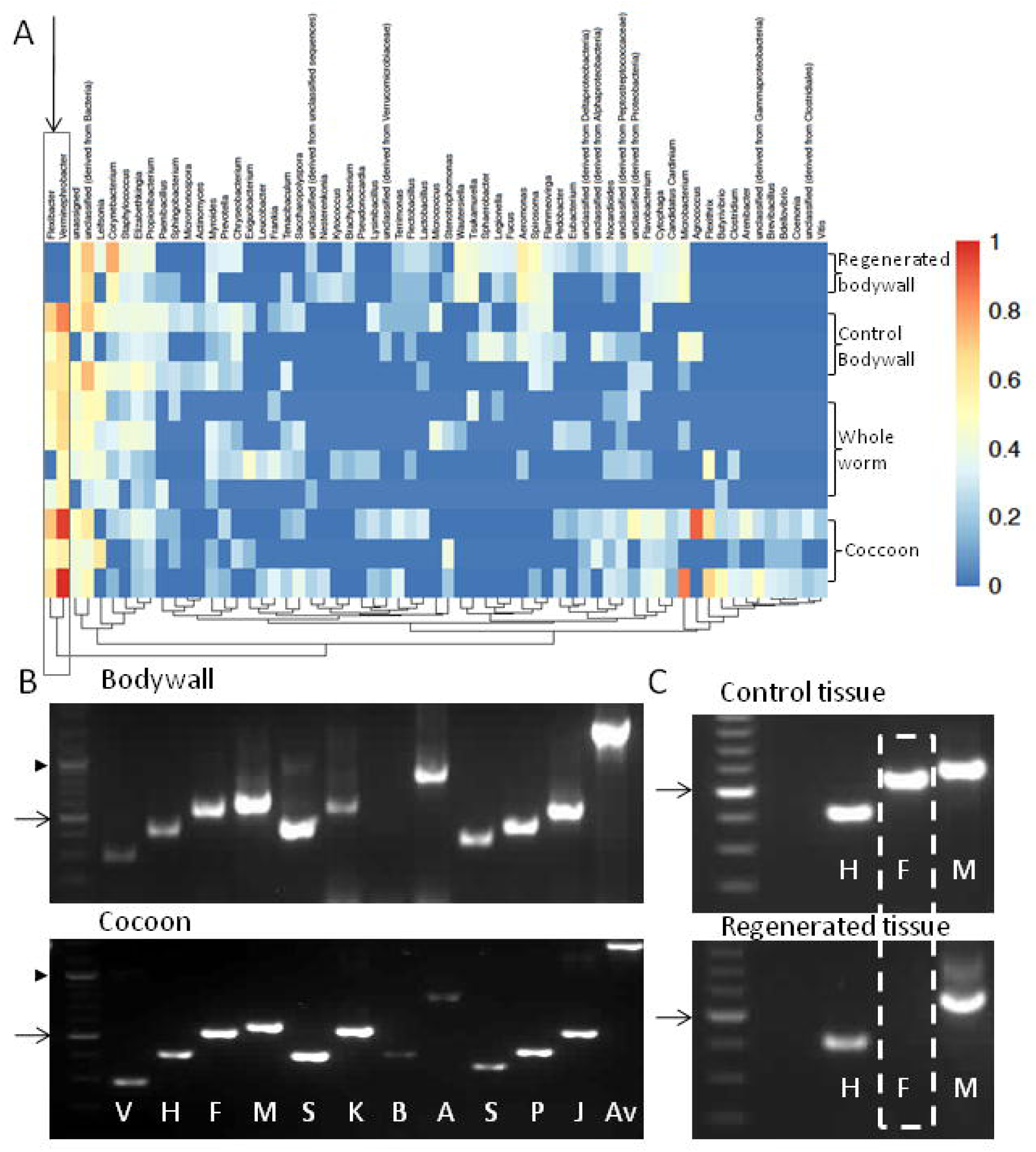
(A) Metagenomics of earthworm tissues by 16S rRNA sequencing of the variable V3 region. Low abundance groups were removed before hierarchical clustering. Scale shows the relative abundance of species. The grey box marked by the arrow shows the two microbial symbionts-*Flexibacter* and *Verminephrobacter* (B) The bodywall and cocoons showed presence of several vertically transmitted microbial species. (C) The regenerated tissue failed to acquire the obligate nephridial symbiont *Flexibacter* species while the adjacent tissue (control) harboured it. DNA isolated from Bodywall, Coccons, Control tissue, or regenerated tissue were subjected to PCR amplification using universal 16s rDNA primers and subsequently re-amplified using specific primers against *Verminephrobacter eiseniae* (V), *Herbaspirillum* (H), *Flexibacter* (F), *Microbacteriaceae* (M), *Sphingomonas* sp. (S), *Klebsiella* sp. (K), *Enterobacter* sp. (E), *Bordetella* sp. (B), *Acidobacteria* (A), *Stenotrophomonas* (S), *Pseudomonas* (P), *Janthinobacterium* (J), *Acidovorax* sp. (Ac) DNA size markers at 1000bp and 500bp are marked with filled arrowheads and open arrows respectively.

Further, we noticed from the metagenome analysis of the regenerated tissue and the adjacent control tissue, that majority of these species were re-instated in the newly grown tissue. Interestingly, the evolutionarily recent symbiont, *Candidatus nephrothrix* was absent in the regenerated tissue, even while it was present in the adjacent tissue (Fig. 7B-C). It appears that the nephridia of the newly regenerated region have no mechanism of acquiring obligate symbionts.

## Discussion

*Eisenia fetida*, or the redworm is a surface dwelling epigeic annelid, distributed on every continent except Antarctica. Genetic mapping and profiling exercises have involved microsatellite markers^77-79^ and AFLP^80^. Attempts have been made at bar-coding species using mitochondrial DNA markers such as COI and COII ^80,81^. Terrestrial annelids are now represented by the genome of *E. fetida* (Indian isolate), reported here and an independently sequenced North American strain ^21^. The ongoing *Lumbricus* genome sequencing effort, is also likely to complement information derived from the *E. fetida* genomes paving the way for a detailed comparative analysis of deep burrowing and surface dwelling worms. For instance, the genome of the highly photophobic *E. fetida* contains several opsin-like sequences. Comparative genomics between deep dwelling and surface dwelling worms in the distribution and wavelength spectra of opsins could reveal interesting insights into molecular basis of behavior.

Studies in comparative regeneration within this group would also provide insight into the gain or loss of regenerative capability over a short evolutionary span. Some species that recently lost regenerative capability might possess a latent ability suppressed due to evolutionary loss of key genes^10^. In Hydra and some vertebrates regeneration has been ‘rescued’ by experimental expression of certain vital signaling molecules, Wnt-signalling pathway members ^82,83^, membrane proteins ^84^ or by suppression of immune responses 85. In annelids, work on marine naidine annelids has yielded that the position and developmental stage of amputations made, can ‘naturally’ rescue regenerative capacities^40^. The ability to regenerate anterior segments has been lost at least 12 times while the posterior regenerative ability was lost at least 4 times during the evolution of annelids ^86^. The genome of *E. fetida* can in future provide the basis for comparative genomics of regenerating and non-regenerating species within the large class of annelids.

Subtractive hybridization experiments in regenerating fragments of *Enchytraeus japonensis* revealed that novel genes (which are not represented in other phyla or even the *Lumbricus rubellus*) were upregulated in regenerative processes hinting towards the involvement of annelid specific genes in regeneration ^87^. Gene expression profiling in anterior regenerating segments of *Perionyx excavatus* also yielded several novel ESTs, involved in regeneration^88^. Transcriptomic analysis of a mixed-stage sample of regeneration and fission in has also been recently attempted ^89^. In our study, we found that 38 amongst the 951 genes upregulated during regeneration had no known orthologue. The large fraction of genes of unknown function suggests that, like in other annelid species, novel factors are involved in *E. fetida* regeneration. Besides these novel factors, well known re-programming genes like SOX4 ^90^ and Brachyury ^52,91^ were also induced in the regenerating tissue. The co-regulation of these and the invertebrate specific factors (conserved differentiation factors) implies that they work in unison. We speculate that novel factors derived from earthworms could participate in cellular networks driven by mammalian reprogramming factors. For instance, our analyses revealed the presence of an earthworm Nerve Growth Factor with only rudimentary similarities to mammalian nerve growth factors, but the Trk receptors showed a higher degree of conservation. In future, functional regeneration assays in mammalian nerve injury models using earthworm derived factors can ascertain the barriers to regeneration in the mammalian nervous system.

Varhalmi et. al. have previously studied the expression of orthologs of pituitary adenylate cyclase-activating polypeptide (PACAP) using radioimmunoassay and immunohistochemistry in *E. fetida* ^92,93^. Since these factors are Neurotrophin in vertebrates, it was thought that their accumulation in the regenerating tissue of *E. fetida* was critical for its innervation. However, the sequences of these factors were not known and the authors relied on immunoreactivity to antibodies against orthologous human proteins. In our experiments, at 15dpc and 20dpc, the earthworm ortholog of PACAP type I receptor was downregulated in the regenerating tissue.

Vertical transmission of metagenome in *E. fetida* was known previously and stands re-affirmed in our study^15^. The inability to restore the obligate symbiont nephridial microbiome post-regeneration is an unexpected finding. Interestingly, while the earthworm seems to regenerate all the other cell types, it fails to acquire the nephridial microbiome.

Earthworms are ubiquitous organisms that impregnate the earth as epigeic surface dwellers, or as endogeic or anecic underground burrowers, ingesting and excreting surface litter or deeper soils, constantly changing its composition and architecture^94^. Estimates of types of earthworms range in thousands, including giant species that are several feet long to others that are only a few centimeters^95^. They also occupy diverse niches, ranging from forest soils to within epiphytes, that live up on trees^6,96^. Their movement due to human activity can threaten entire ecosystems and collectively they can move large amounts of soil, churning a whole ecosystem vertically^1^. They play a vital role in our ecosystems in nutrient cycling, decomposing, vermicomposting and soil aeration. The vast body of literature regarding earthworms bears testimony to their importance as ‘sentinel’ species in environmental and toxicological studies. Biochemical and molecular studies employing proteomics ^97^, transcriptomics ^98,99^ and metabolomics 100 have been attempted to characterize responses to toxins and infectious agents in an effort to exploit them as biomarkers for possible pollutants. Besides their important position in ecology they are also valuable in evolutionary studies as 100 million year old representatives of the versatile and ancient phyllum Annelida that originated during the Cretaceous period^101^.

The emergence of affordable, next generation sequencing technology has enabled comparative genomics of vertebrates, but sequences of invertebrate genomes, especially larger ones has remained scant. By sequencing the genome along with a detailed transcriptomic analysis we were able to identify novel factors involved in invertebrate specific regeneration. Study of invertebrate biology has in the past led to unforeseen applications of human use exemplified by the discovery of Green Fluorescent protein from *Aequorea victoria* that has led to an array of technologies. The earthworm genome and metagenome holds promise of novel factors involved in regeneration and a better understanding of soil, ecology, paving the way for genomics based tools to study genetic diversity. The large diversity of earthworm species suggests that the *E. fetida* genomes can serve as a node for cross-species genome-scale comparisons.

## ACKNOWLEDGEMENT

The authors acknowledge funding support under the EMPOWER grant (OLP1103) from Council for Scientific and Industrial Research, India. Technical assistance in using DESeq2 for differential expression analysis from Megha Lal, IGIB. Technical assistance in histology from Poorti Kathpalia and Department of Anatomy, AIIMS is duly acknowledged. Fellowship from CSIR to Aksheev Bhambri is acknowledged. Fellowship from University Grants Commission(UGC) to Surendra Singh Patel is acknowledged.

### Contributions

Abhishek Bhatt: In-situ hybridization experiments.

Aksheev Bhambri*: Sample collection, library preparation, assembly, regeneration experiments, transcriptome analysis and annotation

Ankit Verma: Genome sequencing

Bastian Fromm: miRNA annotation

Beena Pillai: Conceived the problem, transcriptome analysis, manuscript preparation

Hemant Suryawanshi: Metagenomics

Jameel Ahmed Khan: Nanopore sequencing

Kevin J. Peterson: miRNA annotation

Manish Kumar Rai: Metagenomics sequencing

Mitali Hardikar: Assembly and annotation of RNA-Seq

Nagesh Srikakulam: Genome assembly and annotation

Neeraj Dhaunta*: Sample collection, regeneration experiments, metagenomics

Pradeep Gautam: Histology

Rajesh Pandey: Genome sequencing

Rijith Jayarajan: Genome sequencing

Shamsudheen Vellarikkal: Genome sequencing

Shruti Shridhar: *E. fetida* culture, sequencing data generation, metagenomics, manuscript preparation

Sridhar Sivasubbu: Genome sequencing

Surendra Singh Patel: Transcriptome analysis, manuscript preparation

Vikram Kumar: Genome sequencing

Vinod Scaria: Genome sequencing

## Supplementary data

S1: Primers used in this study

S2: miRNAs annotated from *E. fetida* small RNA transcriptome and genome (separate file)

S3: Analysis of paralogues in *E. fetida* miRNome (separate file)

S4:The mitochondrial genome of *E. fetida*

S5:Annotation of Nerve Growth Factor from *E. fetida*

S4:The mitochondrial genome of *E. fetida*

S5: Nerve Growth Factor from *E. fetida*

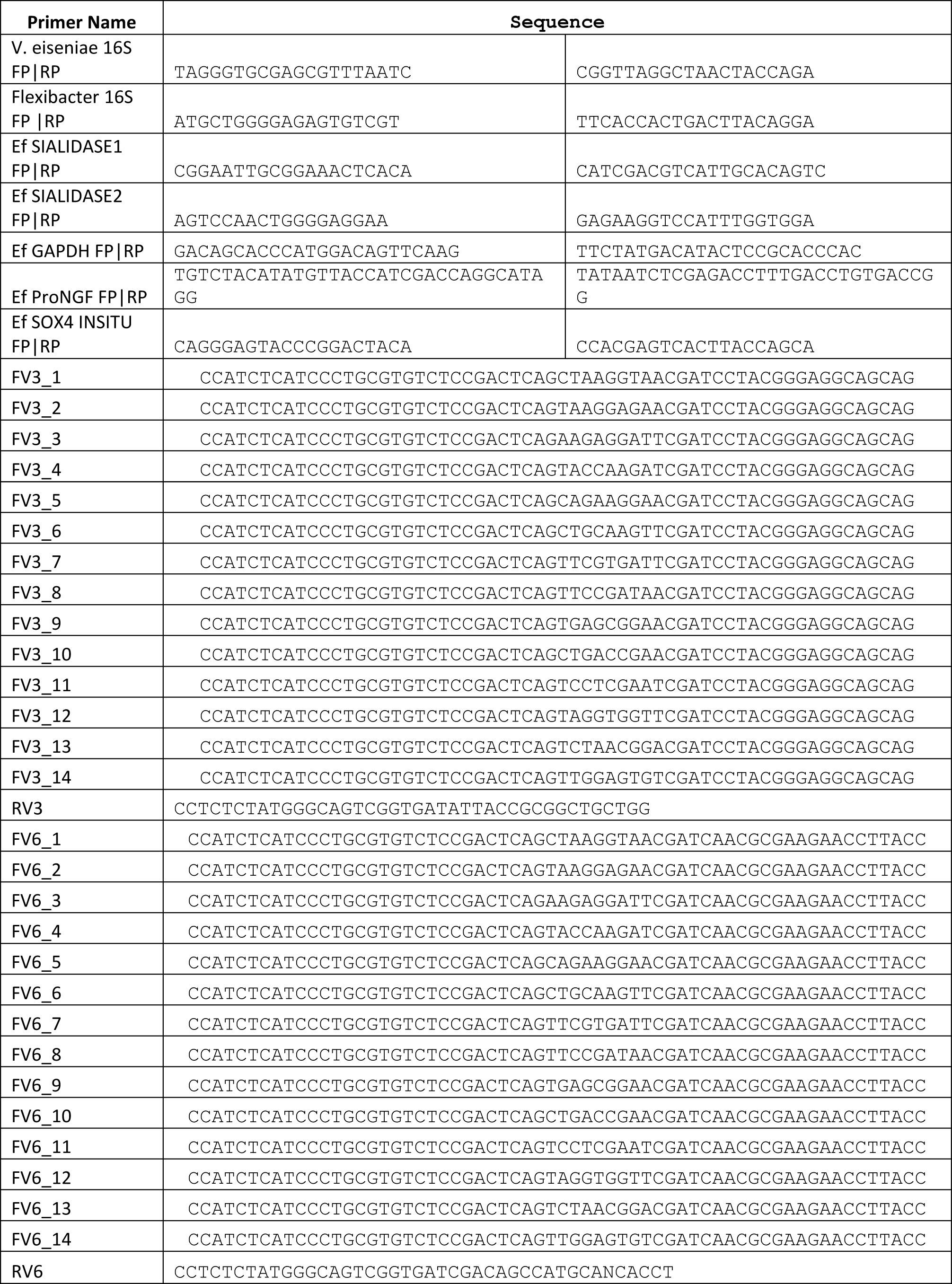

**S4:**
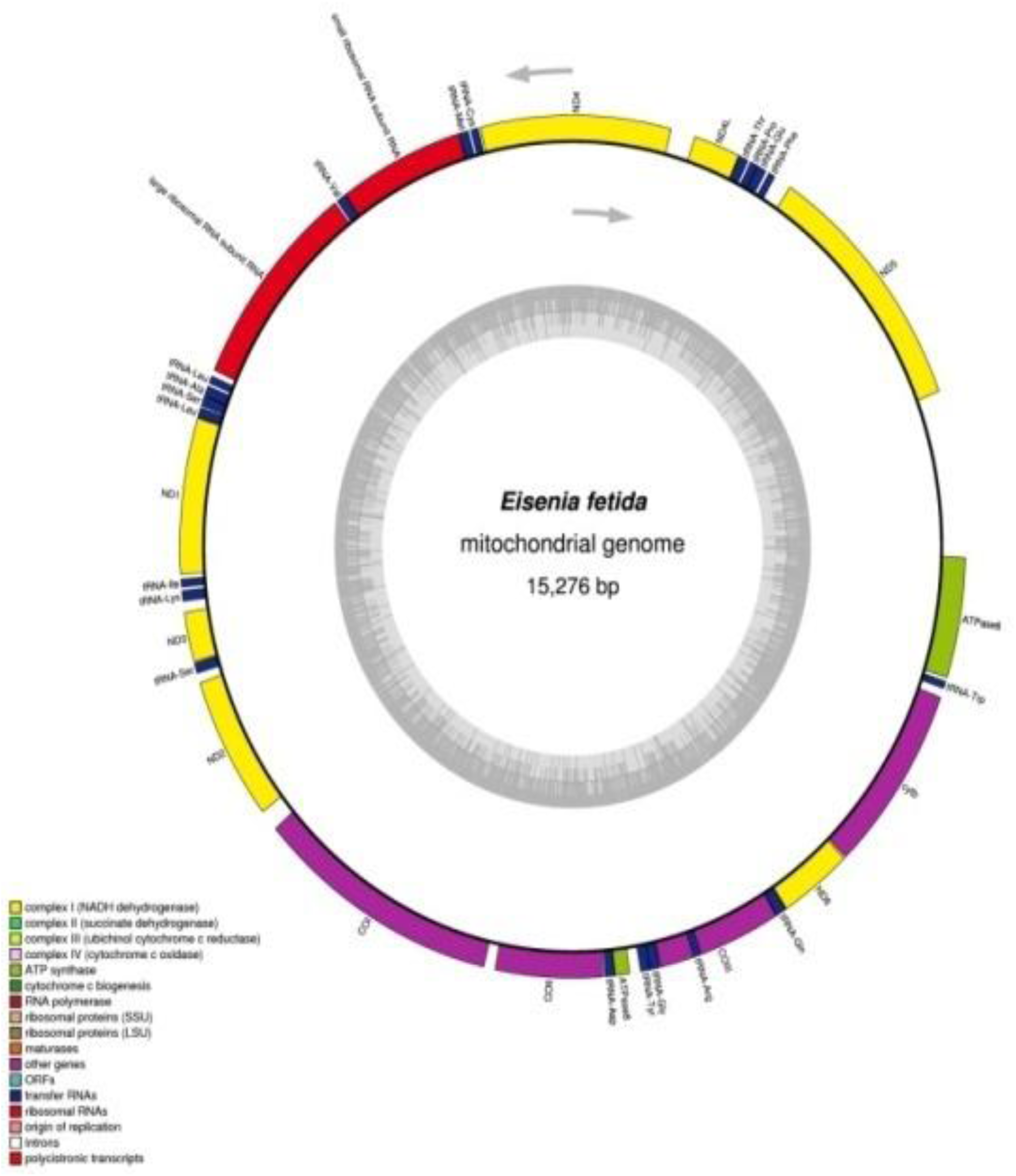
The mitochondrial genome of *E. fetida*

**S5:**
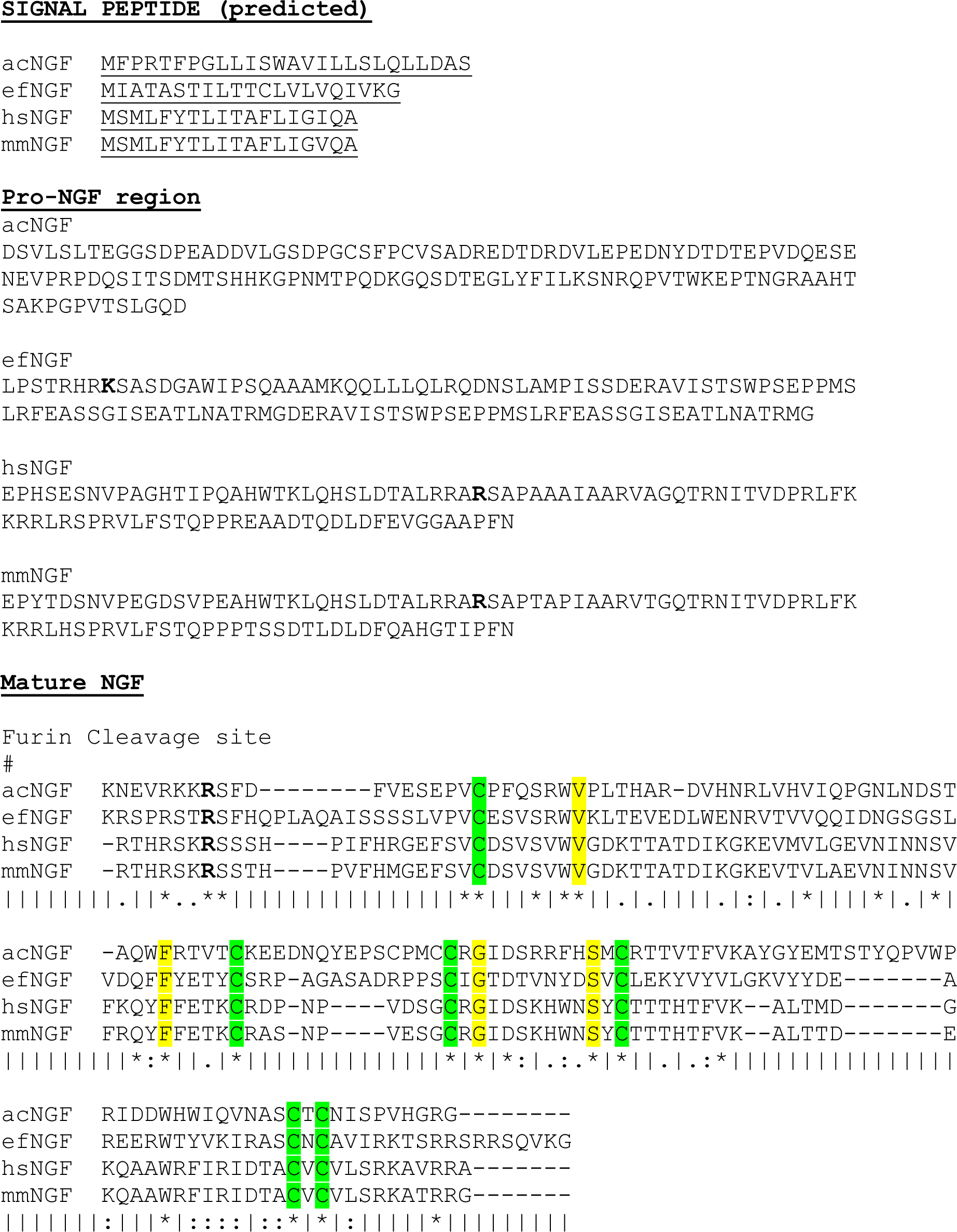
Nerve Growth Factor from *E. fetida*

## Footnotes

* equal contributors

## References

1. Drake, H. L. & Horn, M. A. As the worm turns: the earthworm gut as a transient habitat for soil microbial biomes. Annual review of microbiology 61, 169–189, doi:10.1146/annurev.micro.61.080706.093139 (2007).

2. Uebelacker, J. M. A new parasitic polychaetous annelid (Arabellidae) from the Bahamas. The Journal of Parasitology, 151–154 (1978).

3. Kristensen, E. & Kostka, J. Macrofaunal burrows and irrigation in marine sediment: microbiological and biogeochemical interactions. Interactions between macro- and microorganisms in marine sediments, 125–157 (2005).

4. Parry, L., Vinther, J. & Edgecombe, G. D. Cambrian stem-group annelids and a metameric origin of the annelid head. Biology letters 11 (2015).

5. Brusca, R. C., Brusca, G. J. & Haver, N. J. Invertebrates. (Sinauer Associates Sunderland, Massachusetts, 1990).

6. Edwards, C. A. & Bohlen, P. J. Biology and ecology of earthworms. Vol. 3 (Springer Science & Business Media, 1996).

7. Laumer, C. E. et al. Spiralian phylogeny informs the evolution of microscopic lineages. Current biology : CB 25, 2000–2006, doi:10.1016/j.cub.2015.06.068 (2015).

8. Galliot, B. & Chera, S. The Hydra model: disclosing an apoptosis-driven generator of Wnt-based regeneration. Trends in cell biology 20, 514–523, doi:10.1016/j.tcb.2010.05.006 (2010).

9. Dalyell, J. G. Observations on some interesting phenomena in animal physiology, exhibited by several species of planariae. (Dyer Press, 2007).

10. Bely, A. E. Distribution of segment regeneration ability in the Annelida. Integrative and Comparative Biology 46, 508–518 (2006).

11. Merker, E. & Braunig, G. Die Empfindlichkeit feucchthautiger Tiere im Lichte. 3. Die Empfindlichkeit feucchthautiger Tiere im Licht der Quarzquecksiblerlampe. Zool. Jb. Abt. Allgem. Zool. Physiol, Tiere 43, 275–338 (1927).

12. Moore, M. The rapid escape response of the earthworm Lumbricus terrestris L.: overlapping sensory fields of the median and lateral giant fibres. The Journal of Experimental Biology 83, 231–238 (1979).

13. Mitra, O., Callaham, M. A., Jr., Smith, M. L. & Yack, J. E. Grunting for worms: seismic vibrations cause Diplocardia earthworms to emerge from the soil. Biology letters 5, 16–19, doi:10.1098/rsbl.2008.0456 (2009).

14. Wilson, W. J., Ferrara, N. C., Blaker, A. L. & Giddings, C. E. Escape and avoidance learning in the earthworm Eisenia hortensis. PeerJ 2, e250, doi:10.7717/peerj.250 (2014).

15. Lund, M. B., Kjeldsen, K. U. & Schramm, A. The earthworm-Verminephrobacter symbiosis: an emerging experimental system to study extracellular symbiosis. Frontiers in microbiology 5, 128, doi:10.3389/fmicb.2014.00128 (2014).

16. Consortium, C. e. S. Genome sequence of the nematode C. elegans: a platform for investigating biology. Science 282, 2012–2018 (1998).

17. Adams, M. D. et al. The genome sequence of Drosophila melanogaster. Science 287, 2185–2195 (2000).

18. Howe, K. et al. The zebrafish reference genome sequence and its relationship to the human genome. Nature 496, 498–503, doi:10.1038/nature12111 (2013).

19. Boore, J. L. Complete mitochondrial genome sequence of the polychaete annelid Platynereis dumerilii. Molecular biology and evolution 18, 1413–1416 (2001).

20. Simakov, O. et al. Insights into bilaterian evolution from three spiralian genomes. Nature 493, 526–531, doi:10.1038/nature11696 (2013).

21. Zwarycz, A. S., Nossa, C. W., Putnam, N. H. & Ryan, J. F. Timing and scope of genomic expansion within Annelida: evidence from homeoboxes in the genome of the earthworm Eisenia fetida. Genome biology and evolution, evv243 (2015).

22. el Adlouni, C., Mukhopadhyay, M. J., Walsh, P., Poirier, G. G. & Nadeau, D. Isolation of genomic DNA from the earthworm species Eisenia fetida. Molecular and cellular biochemistry 142, 19–23 (1995).

23. Loman, N. J. & Quinlan, A. R. Poretools: a toolkit for analyzing nanopore sequence data. Bioinformatics 30, 3399–3401 (2014).

24. Meyer, F. et al. The metagenomics RAST server - a public resource for the automatic phylogenetic and functional analysis of metagenomes. BMC bioinformatics 9, 386, doi:10.1186/1471-2105-9-386 (2008).

25. Grabherr, M. G. et al. Full-length transcriptome assembly from RNA-Seq data without a reference genome. Nature biotechnology 29, 644–652, doi:10.1038/nbt.1883 (2011).

26. Langmead, B., Trapnell, C., Pop, M. & Salzberg, S. L. Ultrafast and memory-efficient alignment of short DNA sequences to the human genome. Genome biology 10, R25, doi:10.1186/gb-2009-10-3-r25 (2009).

27. Anders, S., Pyl, P. T. & Huber, W. HTSeq--a Python framework to work with high-throughput sequencing data. Bioinformatics 31, 166–169, doi:10.1093/bioinformatics/btu638 (2015).

28. Love, M. I., Huber, W. & Anders, S. Moderated estimation of fold change and dispersion for RNA-seq data with DESeq2. Genome biology 15, 550, doi:10.1186/s13059-014-0550-8 (2014).

29. Wheeler, B. M. et al. The deep evolution of metazoan microRNAs. Evolution & development 11, 50–68, doi:10.1111/j.1525-142X.2008.00302.x (2009).

30. EA, S. et al. - MicroRNAs resolve an apparent conflict between annelid systematics and their. Proceedings. Biological sciences / The Royal Society 276, 4315–4322 (2009).

31. Folmer, O., Black, M., Hoeh, W., Lutz, R. & Vrijenhoek, R. DNA primers for amplification of mitochondrial cytochrome c oxidase subunit I from diverse metazoan invertebrates. Molecular marine biology and biotechnology 3, 294–299 (1994).

32. Römbke, J. et al. DNA barcoding of earthworms (Eisenia fetida/andrei complex) from 28 ecotoxicological test laboratories. Applied Soil Ecology (2015).

33. Simao, F. A., Waterhouse, R. M., Ioannidis, P., Kriventseva, E. V. & Zdobnov, E. M. BUSCO: assessing genome assembly and annotation completeness with single-copy orthologs. Bioinformatics 31, 3210–3212, doi:10.1093/bioinformatics/btv351 (2015).

34. Boore, J. L. & Brown, W. M. Complete sequence of the mitochondrial DNA of the annelid worm Lumbricus terrestris. Genetics 141, 305–319 (1995).

35. B, F. et al. - A Uniform System for the Annotation of Vertebrate microRNA Genes and the. Annu Rev Genet 49, 213–242 (2015).

36. JE, T. et al. - miRNAs: small genes with big potential in metazoan phylogenetics. Molecular biology and evolution 30, 2369–2382 (2013).

37. A, H. & E, M. - Building a robust a-p axis. Curr Genomics 13, 278–288 (2012).

38. Gates, G. Regeneration in an earthworm, Eisenia foetida (Savigny) 1826. I. Anterior regeneration. The Biological Bulletin 96, 129–139 (1949).

39. Moment, G. B. Segment frequencies in anterior regeneration in the earthworm, Eisenia foetida. The Journal of experimental zoology 111, 449–456 (1949).

40. Bely, A. E. & Sikes, J. M. Latent regeneration abilities persist following recent evolutionary loss in asexual annelids. Proceedings of the National Academy of Sciences of the United States of America 107, 1464–1469, doi:10.1073/pnas.0907931107 (2010).

41. Nengwen, X., Feng, G. & Edwards, C. A. The regeneration capacity of an earthworm, Eisenia fetida, in relation to the site of amputation along the body. Acta Ecologica Sinica (International Journal) 31, 197–204, doi:http://dx.doi.org/10.1016/j.chnaes.2011.04.004 (2011).

42. Clark, F. H. Linkage Studies of Brachyury (Short Tail) in the House Mouse. Proceedings of the National Academy of Sciences of the United States of America 20, 276–279 (1934).

43. Wu, B. et al. Identification of a novel mouse brachyury (T) allele causing a short tail mutation in mice. Cell biochemistry and biophysics 58, 129–135, doi:10.1007/s12013-010-9097-9 (2010).

44. Kispert, A., Herrmann, B. G., Leptin, M. & Reuter, R. Homologs of the mouse Brachyury gene are involved in the specification of posterior terminal structures in Drosophila, Tribolium, and Locusta. Genes & development 8, 2137–2150 (1994).

45. Peterson, K. J., Harada, Y., Cameron, R. A. & Davidson, E. H. Expression pattern of Brachyury and Not in the sea urchin: comparative implications for the origins of mesoderm in the basal deuterostomes. Developmental biology 207, 419–431, doi:10.1006/dbio.1998.9177 (1999).

46. Woollard, A. & Hodgkin, J. The caenorhabditis elegans fate-determining gene mab-9 encodes a T-box protein required to pattern the posterior hindgut. Genes & development 14, 596–603 (2000).

47. Schulte-Merker, S., van Eeden, F. J., Halpern, M. E., Kimmel, C. B. & Nusslein-Volhard, C. no tail (ntl) is the zebrafish homologue of the mouse T (Brachyury) gene. Development 120, 1009–1015 (1994).

48. Benitez, M. S. & Del Pino, E. M. Expression of Brachyury during development of the dendrobatid frog Colostethus machalilla. Developmental dynamics : an official publication of the American Association of Anatomists 225, 592–596, doi:10.1002/dvdy.10190 (2002).

49. Kispert, A., Ortner, H., Cooke, J. & Herrmann, B. G. The chick Brachyury gene: developmental expression pattern and response to axial induction by localized activin. Developmental biology 168, 406–415, doi:10.1006/dbio.1995.1090 (1995).

50. Smith, J. T-box genes: what they do and how they do it. Trends in genetics : TIG 15, 154–158 (1999).

51. Kavka, A. I. & Green, J. B. Tales of tails: Brachyury and the T-box genes. Biochimica et biophysica acta 1333, F73–84 (1997).

52. Technau, U. & Bode, H. R. HyBra1, a Brachyury homologue, acts during head formation in Hydra. Development 126, 999–1010 (1999).

53. Sone, K., Takahashi, T. C., Takabatake, Y., Takeshima, K. & Takabatake, T. Expression of five novel T-box genes and brachyury during embryogenesis, and in developing and regenerating limbs and tails of newts. Development, growth & differentiation 41, 321–333 (1999).

54. Seaver, E. C., Yamaguchi, E., Richards, G. S. & Meyer, N. P. Expression of the pair-rule gene homologs runt, Pax3/7, even-skipped-1 and even-skipped-2 during larval and juvenile development of the polychaete annelid Capitella teleta does not support a role in segmentation. EvoDevo 3, 8, doi:10.1186/2041-9139-3-8 (2012).

55. de Rosa, R., Prud’homme, B. & Balavoine, G. Caudal and even-skipped in the annelid Platynereis dumerilii and the ancestry of posterior growth. Evolution & development 7, 574–587, doi:10.1111/j.1525-142X.2005.05061.x (2005).

56. Brulfert, A., Monnot, M. J. & Geraudie, J. Expression of two even-skipped genes eve1 and evx2during zebrafish fin morphogenesis and their regulation by retinoic acid. The International journal of developmental biology 42, 1117–1124 (1998).

57. Tiwari, N. et al. Sox4 Is a Master Regulator of Epithelial-Mesenchymal Transition by Controlling Ezh2 Expression and Epigenetic Reprogramming. Cancer Cell 23, 768–783, doi:10.1016/j.ccr.2013.04.020.

58. Varki, A. & Schauer, R. Sialic acids. (2009).

59. Takahashi, K. et al. Sialidase NEU4 hydrolyzes polysialic acids of neural cell adhesion molecules and negatively regulates neurite formation by hippocampal neurons. The Journal of biological chemistry 287, 14816–14826, doi:10.1074/jbc.M111.324186 (2012).

60. Varki, A. & Gagneux, P. Multifarious roles of sialic acids in immunity. Annals of the New York Academy of Sciences 1253, 16–36, doi:10.1111/j.1749-6632.2012.06517.x (2012).

61. Surolia, I. et al. Functionally defective germline variants of sialic acid acetylesterase inautoimmunity. Nature 466, 243–247, doi:10.1038/nature09115 (2010).

62. Cariappa, A. et al. B cell antigen receptor signal strength and peripheral B cell development are regulated by a 9-O-acetyl sialic acid esterase. The Journal of experimental medicine 206, 125–138, doi:10.1084/jem.20081399 (2009).

63. Miyagi, T. & Yamaguchi, K. Mammalian sialidases: physiological and pathological roles in cellular functions. Glycobiology 22, 880–896, doi:10.1093/glycob/cws057 (2012).

64. Dvorak, J. et al. Microbial environment affects innate immunity in two closely related earthworm species Eisenia andrei and Eisenia fetida. PloS one 8, e79257, doi:10.1371/journal.pone.0079257 (2013).

65. McLean, K. E. & Vickaryous, M. K. A novel amniote model of epimorphic regeneration: the leopard gecko, Eublepharis macularius. BMC developmental biology 11, 50, doi:10.1186/1471-213X-11-50 (2011).

66. Myers, M. W., Lazzarini, R. A., Lee, V. M., Schlaepfer, W. W. & Nelson, D. L. The human mid-size neurofilament subunit: a repeated protein sequence and the relationship of its gene to the intermediate filament gene family. The EMBO journal 6, 1617–1626 (1987).

67. Yuan, A., Rao, M. V. & Nixon, R. A. Neurofilaments at a glance. Journal of cell science 125, 3257–3263 (2012).

68. Wang, H. et al. Neurofilament proteins in axonal regeneration and neurodegenerative diseases. Neural regeneration research 7, 620–626, doi:10.3969/j.issn.1673-5374.2012.08.010 (2012).

69. Wallwork, J. A. Earthworm biology. (E. Arnold (Publishers) Ltd., 1983).

70. Levi, J. U., Cowden, R. R. & Collins, G. H. The microscopic anatomy and ulrastructure of thenervous system in the earthworm (Lumbricus sp.) with emphasis on the relationship between glial cells and neurons. Journal of Comparative Neurology 127, 489–509 (1966).

71. Painter, B. T. The location of factors of head regeneration in the earthworm. The Biological Bulletin 78, 463–485 (1940).

72. Taniuchi, M., Clark, H. B., Schweitzer, J. B. & Johnson, E. M., Jr. Expression of nerve growthfactor receptors by Schwann cells of axotomized peripheral nerves: ultrastructural location,suppression by axonal contact, and binding properties. The Journal of neuroscience : the official journal of the Society for Neuroscience 8, 664–681 (1988).

73. Kassabov, S. R. et al. A single Aplysia neurotrophin mediates synaptic facilitation via differentially processed isoforms. Cell reports 3, 1213–1227, doi:10.1016/j.celrep.2013.03.008(2013).

74. Valembois, P., Lassegues, M. & Roch, P. Formation of brown bodies in the coelomic cavity of the earthworm Eisenia fetida andrei and attendant changes in shape and adhesive capacity of constitutive cells. Developmental and comparative immunology 16, 95–101 (1992).

75. Engelmann, P., Molnár, L., Pálinkás, L., Cooper, E. & Németh, P. Earthworm leukocyte populations specifically harbor lysosomal enzymes that may respond to bacterial challenge. Cell and tissue research 316, 391–401 (2004).

76. Pinel, N., Davidson, S. K. & Stahl, D. A. Verminephrobacter eiseniae gen. nov., sp. nov., anephridial symbiont of the earthworm Eisenia foetida (Savigny). International journal of systematic and evolutionary microbiology 58, 2147–2157, doi:10.1099/ijs.0.65174-0 (2008).

77. Huang, J., Xu, Q., Sun, Z. J., Tang, G. L. & Su, Z. Y. Identifying earthworms through DNA barcodes. Pedobiologia 51, 301–309 (2007).

78. Harper, G. L. et al. Microsatellite markers for the earthworm Lumbricus rubellus. Molecular Ecology Notes 6, 325–327 (2006).

79. Velavan, T., Schulenburg, H. & Michiels, N. K. Development and characterization of novel microsatellite markers for the common earthworm (Lumbricus terrestris L.). Molecular Ecology Notes 7, 1060–1062 (2007).

80. King, R. A., Tibble, A. L. & Symondson, W. O. Opening a can of worms: unprecedented sympatric cryptic diversity within British lumbricid earthworms. Molecular ecology 17, 4684–4698, doi:10.1111/j.1365-294X.2008.03931.x (2008).

81. Heethoff, M., Etzold, K. & Scheu, S. Mitochondrial COII sequences indicate that the parthenogenetic earthworm Octolasion tyrtaeum (Savigny 1826) constitutes of two lineages differing in body size and genotype. Pedobiologia 48, 9–13 (2004).

82. Kawakami, Y. et al. Wnt/beta-catenin signaling regulates vertebrate limb regeneration. Genes & development 20, 3232–3237, doi:10.1101/gad.1475106 (2006).

83. Lengfeld, T. et al. Multiple Wnts are involved in Hydra organizer formation and regeneration. Developmental biology 330, 186–199, doi:10.1016/j.ydbio.2009.02.004 (2009).

84. Adams, D. S., Masi, A. & Levin, M. H+ pump-dependent changes in membrane voltage are an early mechanism necessary and sufficient to induce Xenopus tail regeneration. Development 134, 1323–1335, doi:10.1242/dev.02812 (2007).

85. Fukazawa, T., Naora, Y., Kunieda, T. & Kubo, T. Suppression of the immune response potentiates tadpole tail regeneration during the refractory period. Development 136, 2323–2327, doi:10.1242/dev.033985 (2009).

86. Martinez-Acosta, V. G. & Zoran, M. J. Evolutionary Aspects of Annelid Regeneration. eLS.

87. Myohara, M., Niva, C. C. & Lee, J. M. Molecular approach to annelid regeneration: cDNA subtraction cloning reveals various novel genes that are upregulated during the large-scale regeneration of the oligochaete, Enchytraeus japonensis. Developmental dynamics : an official publication of the American Association of Anatomists 235, 2051–2070, doi:10.1002/dvdy.20849(2006).

88. Cho, S.-J. et al. Gene expression profile in the anterior regeneration of the earthworm using expressed sequence tags. Bioscience, biotechnology, and biochemistry 73, 29–34 (2009).

89. Nyberg, K. G., Conte, M. A., Kostyun, J. L., Forde, A. & Bely, A. E. Transcriptome characterization via 454 pyrosequencing of the annelid Pristina leidyi, an emerging model for studying the evolution of regeneration. BMC genomics 13, 287, doi:10.1186/1471-2164-13-287 (2012).

90. Foronda, M. et al. Sox4 links tumor suppression to accelerated aging in mice by modulating stemcell activation. Cell reports 8, 487–500, doi:10.1016/j.celrep.2014.06.031 (2014).

91. Vidricaire, G., Jardine, K. & McBurney, M. W. Expression of the Brachyury gene during mesoderm development in differentiating embryonal carcinoma cell cultures. Development 120, 115–122 (1994).

92. Boros, A. et al. Changes in the expression of PACAP-like compounds during the embryonic development of the earthworm Eisenia fetida. Journal of molecular neuroscience : MN 36, 157–165, doi:10.1007/s12031-008-9102-6 (2008).

93. Varhalmi, E. et al. Expression of PACAP-like compounds during the caudal regeneration of the earthworm Eisenia fetida. Journal of molecular neuroscience : MN 36, 166–174, doi:10.1007/s12031-008-9125-z (2008).

94. Moore, J. & Overhill, R. An introduction to the invertebrates. (Cambridge University Press, 2001).

95. Edwards, C. A. Earthworm ecology. (CRC Press, 2004).

96. Fragoso, C. & Lavelle, P. Earthworm communities of tropical rain forests. Soil Biology and Biochemistry 24, 1397–1408 (1992).

97. Ji, C. et al. Proteomic and metabolomic analysis of earthworm Eisenia fetida exposed to different concentrations of 2,2’,4,4’-tetrabromodiphenyl ether. Journal of proteomics 91, 405–416, doi:10.1016/j.jprot.2013.08.004 (2013).

98. Gong, P. et al. Transcriptomic analysis of RDX and TNT interactive sublethal effects in the earthworm Eisenia fetida. BMC genomics 9 Suppl 1, S15, doi:10.1186/1471-2164-9-S1-S15(2008).

99. Gong, P. et al. Toxicogenomic analysis provides new insights into molecular mechanisms of the sublethal toxicity of 2,4,6-trinitrotoluene in Eisenia fetida. Environmental science & technology 41, 8195–8202 (2007).

100. Bundy, J. G. et al. ’Systems toxicology’ approach identifies coordinated metabolic responses to copper in a terrestrial non-model invertebrate, the earthworm Lumbricus rubellus. BMC biology 6, 25, doi:10.1186/1741-7007-6-25 (2008).

101. Parry, L., Tanner, A. & Vinther, J. The origin of annelids. Palaeontology 57, 1091–1103 (2014).

